# High-fat diet-induced activation of SGK1 promotes Alzheimer’s disease-associated tau pathology

**DOI:** 10.1101/2020.05.14.095471

**Authors:** Montasir Elahi, Yumiko Motoi, Shotaro Shimonaka, Yoko Ishida, Hiroyuki Hioki, Masashi Takanashi, Koichi Ishiguro, Yuzuru Imai, Nobutaka Hattori

**Affiliations:** Department of Diagnosis, Prevention and Treatment of Dementia, Tokyo 113-8421, Japan; Department of Neurology, Juntendo University Graduate of Medicine, Tokyo 113-8421, Japan; Department of Cell Biology and Neuroscience, Juntendo University Graduate of Medicine, Tokyo 113-8421, Japan; Department of Research for Parkinson’s Disease, Juntendo University Graduate School of Medicine, Tokyo 113-8421, Japan

## Abstract

Type2 diabetes mellitus (T2DM) has long been considered a risk factor for Alzheimer’s disease (AD). However, the molecular links between T2DM and AD remain obscure. Here, we reported that serum/glucocorticoid-regulated kinase1 (SGK1) is activated by administering a chronic high-fat diet (HFD), which increases the risk of T2DM, and thus promotes Tau pathology via the phosphorylation of tau at Ser214 and the activation of a key tau kinase, namely, GSK-3ß, forming SGK1-GSK-3ß-tau complex. SGK1 was activated under conditions of elevated glucocorticoid and hyperglycemia associated with HFD, but not of fatty acid-mediated insulin resistance. Elevated expression of SGK1 in the mouse hippocampus led to neurodegeneration and impairments in learning and memory. Upregulation and activation of SGK1, SGK1-GSK-3ß-tau complex were also observed in the hippocampi of AD cases. Our results suggest that SGK1 is a key modifier of tau pathology in AD, linking AD to corticosteroid effects and T2DM.

## Introduction

Alzheimer’s disease (AD) affects more than 45 million people worldwide and is characterized by the accumulation of neurofibrillary tangles (NFTs) composed of hyperphosphorylated tau protein and amyloid plaques primarily composed of aggregated Aβ in affected brain regions such as the hippocampus (GBD 2016 Dementia Collaborators, 2019; Grundke-Iqbal et al., 1986). Although the etiology of AD is not yet fully understood, epidemiological studies have revealed that type 2 diabetes mellitus (T2DM) is a risk factor for AD, as those with T2DM have a 2.5-fold increased risk for experiencing cognitive dysfunction compared to aged-matched individuals (Biessels et al., 2006; Iqbal et al., 2014; Spauwen et al., 2013). Microtubule-associated protein tau, which stabilizes microtubules in neurons, is subject to various posttranslational modifications. Phosphorylation plays a major role in microtubule dysfunction. Tau can be phosphorylated at multiple sites in response to activation of various signaling pathways, including those implicated in metabolic signaling (Grundke-Iqbal et al., 1986; Hanger et al., 2014; Iqbal et al., 2009; Stieler et al., 2011). Phospho-modification is closely linked to tau oligomerization and insolubility (Noble et al., 2013). Indeed, AD patients who also had T2DM have presented with prominent tau pathology in their hippocampi (Valente et al., 2010). Collaterally, diabetes is also suggested to be associated with an increased risk for progression from mild cognitive impairment to dementia (Livingston et al., 2017).

Elevated levels of hyperphosphorylated tau and cognitive and memory impairments have been documented in high-fat diet (HFD)-treated T2DM mice and in leptin receptor-deficient *db/db* mice, a widely used genetic model of T2DM (Johnson et al., 2016; Koga et al., 2014; Leboucher et al., 2013; Li et al., 2002). In HFD-treated obese mice, increased tau phosphorylation associated with tau pathology was often assumed to be linked to dysregulation of the central and peripheral insulin signaling pathway (Bhat and Thirumangalakudi, 2013; Peng et al., 2013; Platt et al., 2016; Sajan et al., 2016). However, elevated tau phosphorylation at various sites was observed regardless of insulin signaling activity (Leboucher et al., 2013; Takalo et al., 2014). Glycogen synthase kinase-3ß (GSK-3ß) is a key tau kinase that promotes tau pathology and is shown to phosphorylate tau at multiple sites (Hanger and Noble, 2011; Ishiguro et al., 1995). However, activation of insulin/phosphatidylinositol-3 kinase(PI3K)/AKT signaling by an HFD negatively regulates GSK-3ß through Ser9 phosphorylation (Leboucher et al., 2013; Sajan et al., 2016), while insulin resistance diminishes AKT activity, thus leading to GSK-3ß activation. Kinases other than GSK-3ß that are shown to affect tau phosphorylation, such as cyclin-dependent kinase 5 (CDK5), mitogen-activated protein kinase (MAPK), Ca^2+^/calmodulin-dependent kinase II (CaMKII), protein kinase C (PKC) and protein phosphatase 2A, remained unchanged in HFD-treated mouse brains (Leboucher et al., 2013; Sajan et al., 2016). In contrast, elevated corticosteroids led to increased activation of GSK-3ß in *db/db* mice though an unknown mechanism (Dey et al., 2017). Therefore, an in-depth analysis of the tau kinases activated by the metabolic changes in diabetes is necessary to understand the molecular mechanism by which altered energy metabolism affects tau stability and subsequent AD development.

Using microarray analysis, we found that HFD induced upregulation and activation of SGK1, which stimulated tau phosphorylation at Ser214 in the mouse hippocampus (Ackermann et al., 2011; Yang et al., 2006). Overexpression of SGK1 in the mouse hippocampus resulted in neurodegeneration and impaired the cognitive functions. Treatment of cultured SH-SY5Y cells with dexamethasone or high glucose activated SGK1 while palmitic acid treatment, which mimics insulin resistance, did not have the same effects, suggesting that elevated glucocorticoid and hyperglycemia were associated with SGK1-mediated tau pathology. Moreover, administration of SGK1 specific inhibitor EMD638683 suppressed the elevated tau phosphorylation in stable tau expressing SH-SY5Y cells and tauopathy model mice, suggesting that SGK1 is a promising therapeutic target in the prodromal stage of AD.

## Results

### HFD administration upregulates SGK1

Tg601 mice neuronally expressing the 2N4R form of human tau develop memory impairments at 17 months of age (Kambe et al., 2011). Tg601 mice and non-transgenic (NTg) mice began on an HFD or normal diet at 10 months of age, when tau pathology and memory impairment are still absent in Tg601 mice. Although Tg601 mice exhibited lower body weight than NTg mice, both Tg601 and NTg mice developed obesity and high fasting glycemia caused by HFD when given a chronic HFD treatment for 5 months (Figure 1—figure supplement 1A and B). Moreover, the intraperitoneal glucose tolerance test (IPGTT) and insulin tolerance test (ITT) indicated that both Tg601 and NTg mice exhibited impaired glucose tolerance and a reduced response to insulin (Figure 1—figure supplement 1C-F).

After establishing human tau transgenic mice experiencing symptoms of T2DM (Tg601 with HFD), we next examined the consequence of increased tau expression and hyperglycemia on hippocampal functions. Previous studies in HFD rodent models have shown that the hippocampus is the most affected part of the brain in diabetes (Elahi et al., 2016a; Stranahan et al., 2008). Behavioral analyses using the Morris water maze, Y-maze spontaneous alteration task and elevated plus maze test revealed that HFD treatment caused impairments in spatial learning, memory function and exploratory behavior and caused anxiety-like behavior to develop in both Tg601 and NTg mice compared with mice that were given a normal diet treatment (Figure 1—figure supplement 2A, D-F). In contrast, swimming speed in the Morris water maze and rotarod performance were similar among all groups, suggesting that motor function was not affected by HFD (Figure 1—figure supplement 2B, C and G). These behavioral tests indicated that HFD treatment accelerated the defects in hippocampal function regardless of tau expression levels.

We next searched for molecules that modulate tau pathology by HFD. Gene expression microarray showed that only a few genes were transcriptionally altered in the hippocampi of HFD-treated mice (Figure 1A). Among the altered genes, *sgk1* showed the highest degree of change (4.0-fold increase) in both Tg601 and NTg mice after HFD treatment (Figure 1A and B). Increased *sgk1* expression at the transcriptional and post-transcriptional levels was verified by RT-PCR and Western blot, respectively (Figure 1C and D). Moreover, phosphorylation of SGK1 at Thr256 and Ser422 was elevated, which involves insulin signaling-associated PDK1 kinase (Chen et al., 2009). These findings suggest that SGK1 kinase activity was increased by the phosphorylation (Figure 1D) (Park et al., 1999). Pronounced phosphorylation of IRS1 at Ser632 (Ser636 in humans) and of AKT at Ser473 was also observed, indicating that insulin signaling was overactivated by HFD (Figure 1D). Consistent with the above observations, SGK1 activity was increased in the hippocampi of HFD-treated mice regardless of tau expression levels (Figure 1E) (Murray et al., 2005). HFD treatment caused an increase in glucocorticoid levels, which has been reported to upregulate SGK1 transcriptionally and causes dysregulation of insulin signaling in mice (Figure 1F) (Hinds et al., 2017). Thus, these results suggest that elevated glucocorticoids and altered insulin signaling in the hippocampi of HFD-treated mice cause both upregulated expression and increased activation of SGK1.

**Figure 1.**
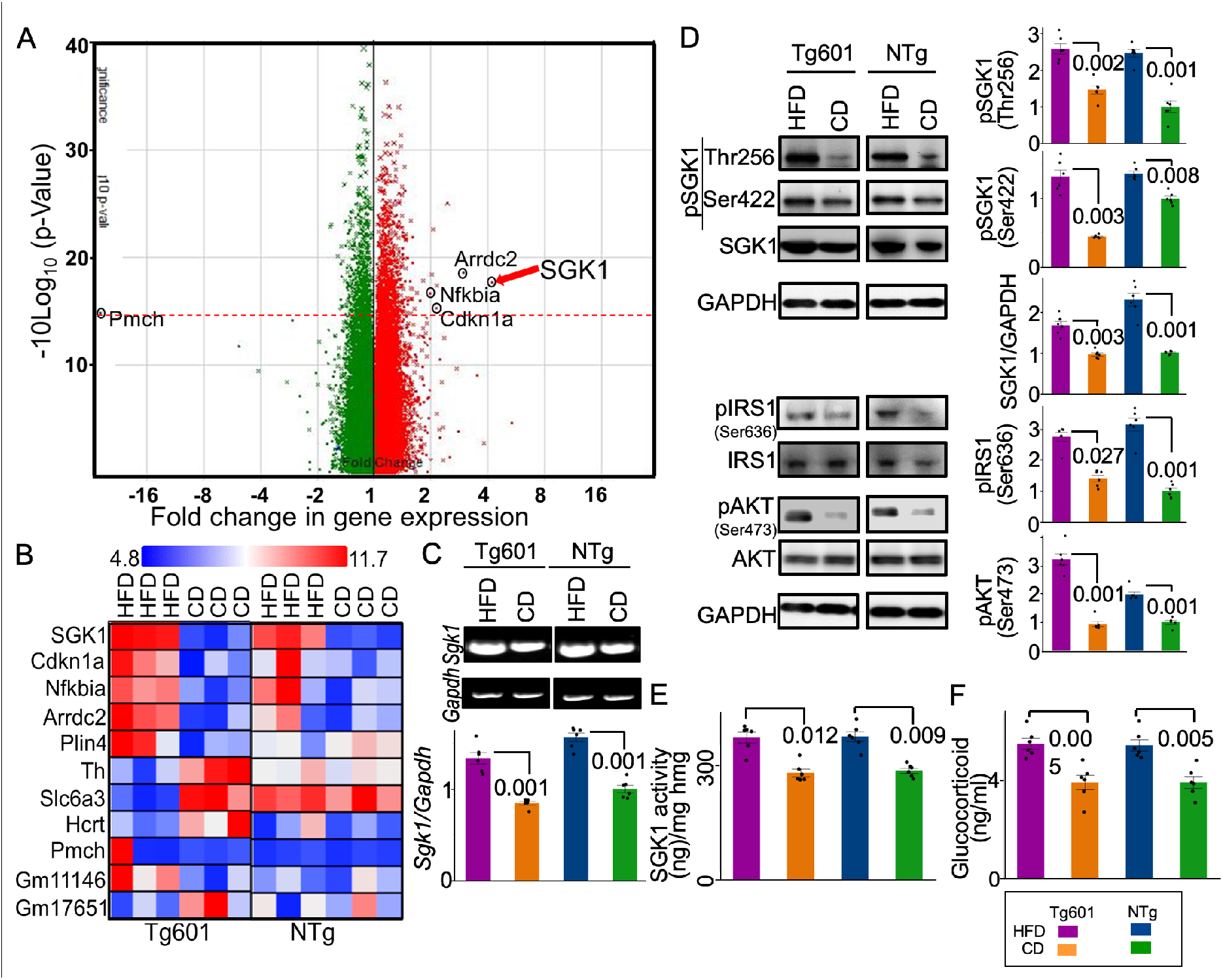
Elevated SGK1 expression and activation in the HFD-treated mouse hippocampus. **(A)** Alterations in gene expression in the HFD-treated Tg601 hippocampus (n = 6 mice) (green, negatively regulated genes; red, positively regulated genes). A broken red line represents *p* = 0.05 and significantly changed genes are indicated. **(B)** A comparative cell plot view of the hippocampal genes with ≥ 2-fold change. **(C)** Validation of the transcriptome array result by RT-PCR (n = 6 biological replicates). The relative values normalized to *Gapdh* were graphed. The value of NTg with CD was set as 1. **(D)** Quantitative Western blotting analysis for the indicated proteins in the hippocampus. Total SGK1 and phosphorylated SGK1 intensities were standardized to GAPDH and total SGK1, respectively (n = 6 mice). **(E)** SGK1 kinase activity using SGK1 specific peptide (SGKtide) as a substrate (n = 6 biological replicates). **(F)** Glucocorticoid levels in mouse hippocampal homogenates (n = 6 mice). Graphs in **C-F** represent means ± SEM and *p* values are indicated in the graphs. Two-way ANOVA followed by Tukey-HSD (**C-E**) and Mann-Whitney U test (**F**).

**Figure 1-figure supplement 1.**
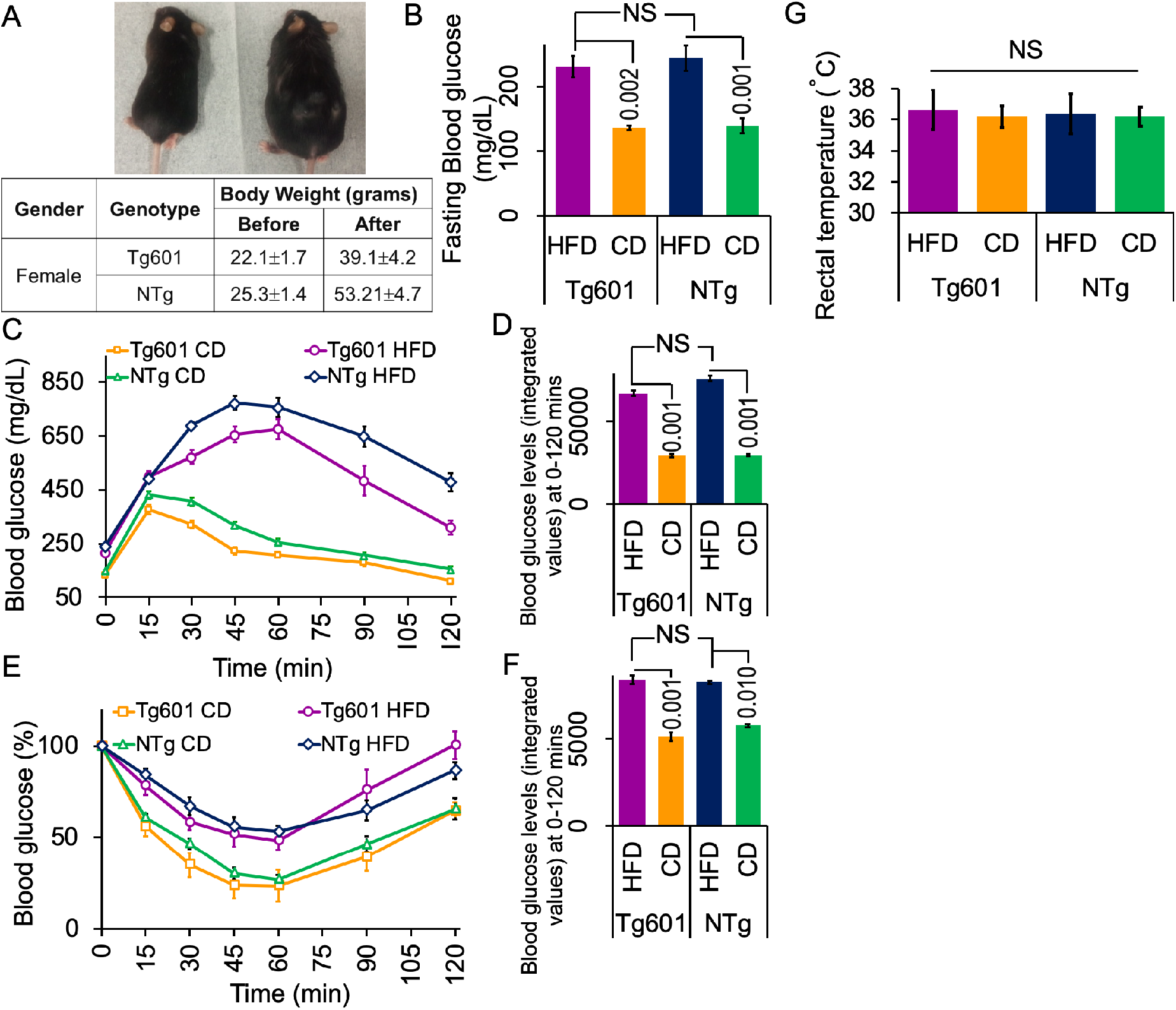
HFD-treated mice exhibit obesity, glucose intolerance and peripheral insulin resistance. **(A)** Appearance of HFD-treated (right) and CD-treated (left) Tg601 mice (upper). Body weights of mice with the indicated genotypes after five months of HFD treatment (lower). **(B)** HFD treatment elevates fasting blood glucose levels. Fasting blood glucose was measured 5 months after the diet treatment, allowing the mice to fast for 8 hrs prior to measurement. **(C)** HFD impairs the control of glucose load. An intraperitoneal glucose tolerance test was performed using HFD or CD mice with the indicated genotypes for the indicated periods of time. **(D)** A graph represents the integrated values of blood glucose levels from 0 to 120 min as in **(C)**. **(E)** HFD evokes insulin resistance. An intraperitoneal insulin tolerance test was performed using HFD or CD mice with the indicated genotypes. **(F)** A graph represents the integrated values of blood glucose levels from 0 to 120 min as in **(E)**. **(G)** The body temperature is not different among the mouse groups. Graph indicated rectal temperature records. Data are represented as the means ± SEM (n = 27 mice in each group) and were analyzed by two-way ANOVA followed by Tukey HSD. *p*-values are indicated in the graphs when significantly different. NS, not significant.

**Figure 1-figure supplement 2.**
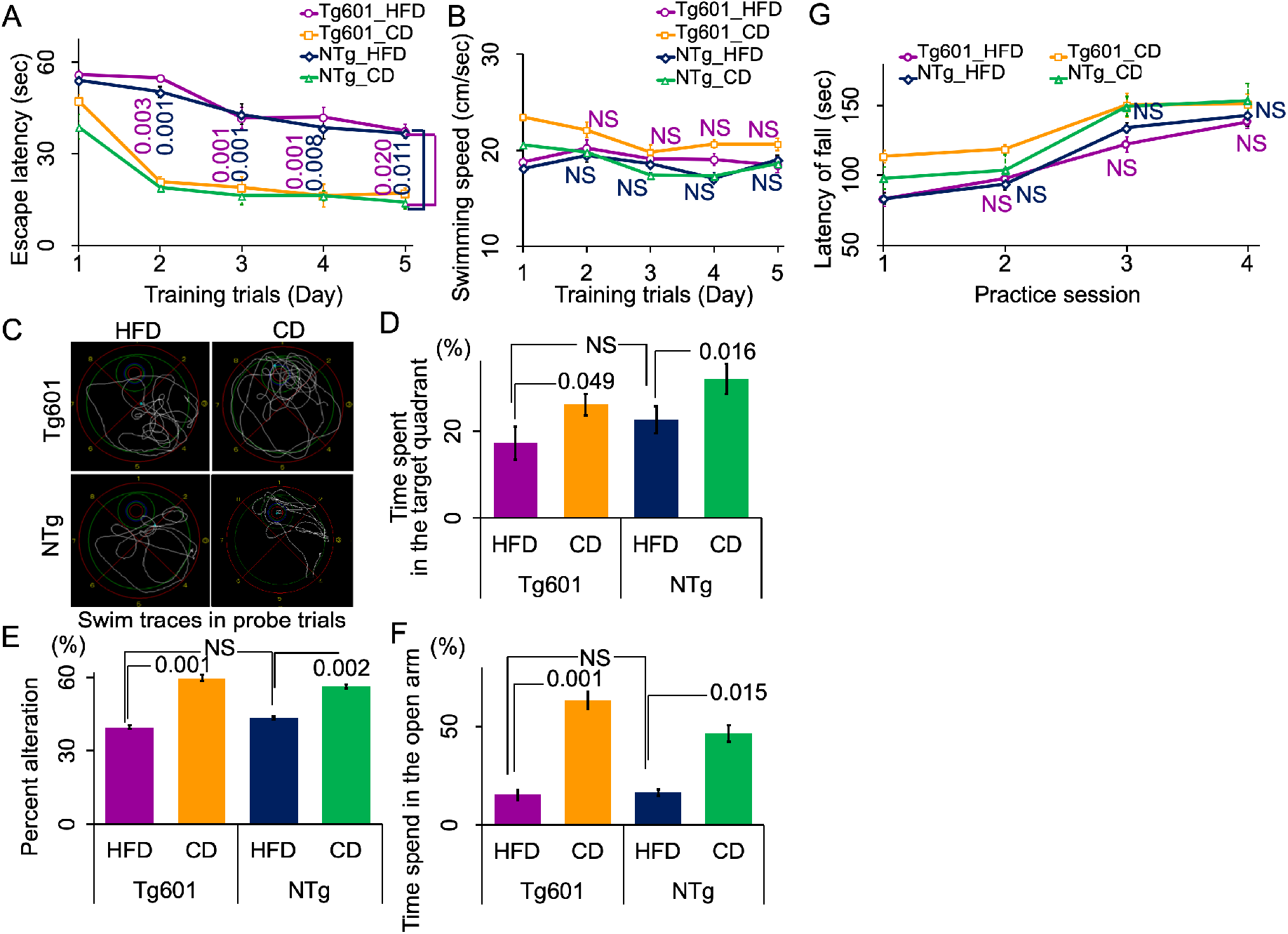
HFD impairs learning and memory and increases anxiety. **(A)** HFD worsens spatial learning. Hidden platform trials were performed in the Morris water maze. HFD mice had significantly longer escape latencies than CD mice. **(B)** Swimming speed in training trials showed no difference in motor function in each group. **(C)** Representative swim traces for each group in probe trials of the Morris water maze test. **(D)** HFD worsens memory function. Percent of goal quadrant dwell time in the probe trials. Probe trials were conducted 48 hrs after the last training session for 3 trials. **(E)** HFD worsens spatial memory. Percent of spontaneous alternations by HFD and CD mice in the Y maze test. HFD treatment reduced spontaneous alternations of Tg601 and NTg mice. **(F)** Anxiety test. Fear feelings were assessed by the time spent in open arms in an elevated plus-maze test. The HFD group showed behavior that is indicative of increased fear feelings. **(G)** HFD does not affect motor function. Motor activity was analyzed by rotarod performance test. Data are represented as the means ± SEM (n = 27 mice in each group) and were analyzed by two-way ANOVA followed by Tukey HSD. *p*-values are indicated in the graphs when significantly different. NS, not significant. **Figure 1—source dataset 1** Source dataset for Figure 1A (not included in this review)

### SGK1 upregulation and activation leads to impaired cognitive functions

Since SGK1 was reported to phosphorylate tau at Ser214, we analyzed the levels of tau phosphorylation at Ser214 (pSer214-phospho-tau) in the hippocampi of HFD-treated mice (Virdee et al., 2007; Yang et al., 2006). Increased levels of Ser214-phospho-tau were found when assessing both transgenic human tau and endogenous murine tau in the hippocampi of Tg601 and NTg mice that were treated with HFD (Figure 2A-D and Figure 2—figure supplement 1A).

**Figure 2.**
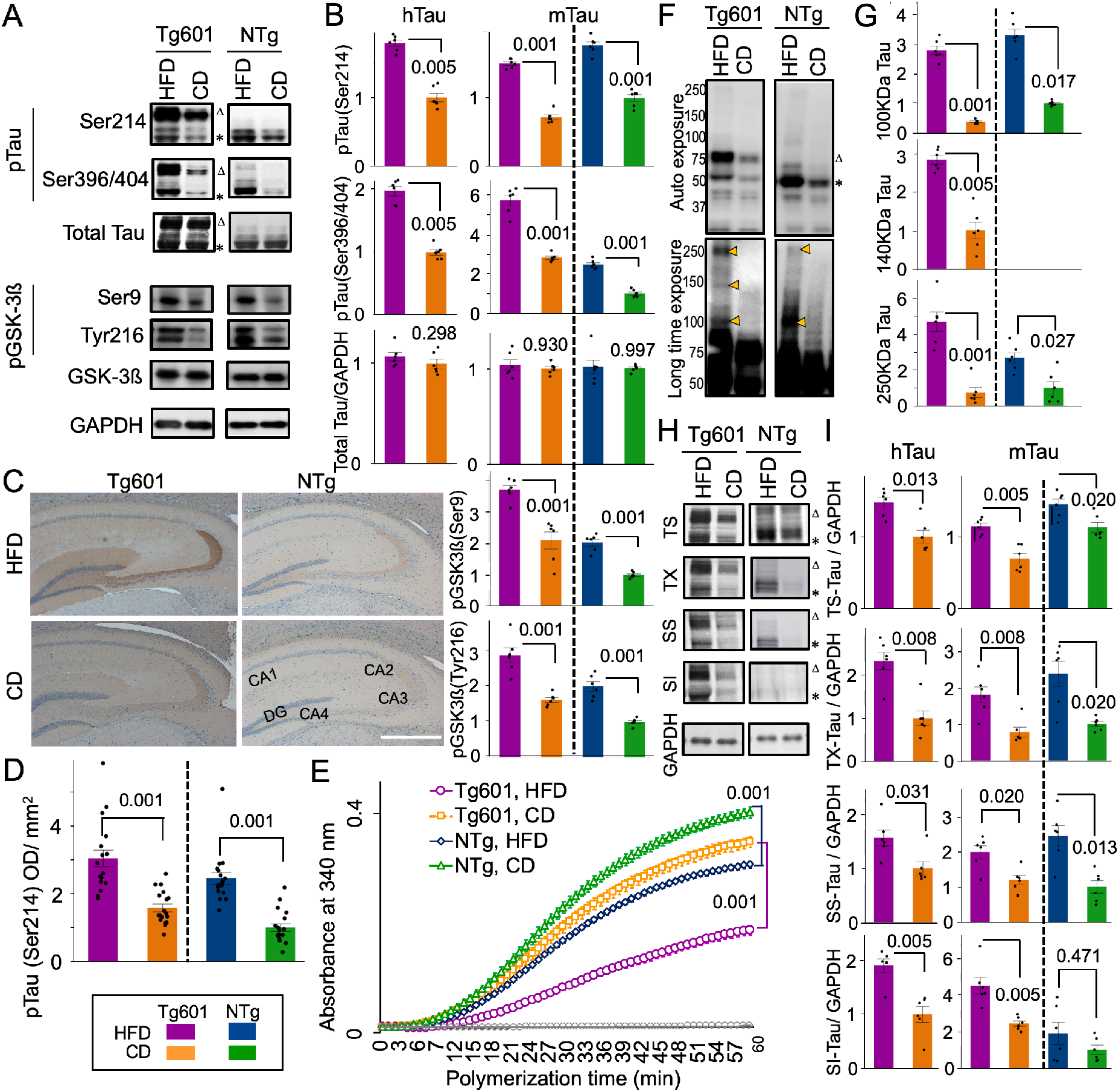
Enhanced Ser214 tau phosphorylation and tau pathology in HFD-treated mice. **(A** and **B)** Quantitative Western blot for the indicated proteins in the mouse brain hippocampus. Transgenic human (Δ: ~75 kDa, hTau) and endogenous murine (*: ~50 kDa, mTau) tau were indicated. Graphs represent means ± SEM (n = 6 mice, arbitrary units). Each control is set to 1. **(C** and **D)** HFD promotes tau pSer214 phosphorylation in the hippocampus. **(C)** Immunohistochemical analysis of pSer214 tau (brown) in HFD- or CD-treated hippocampus counterstained with hematoxylin (blue). Strong pSer214 signals were observed in the mossy fibers in CA3 and neuropils in CA4 in Tg601 mice. DG, dentate gyrus. (**D)** Quantification of pSer214 immunoreactivity in the entire hippocampus region as in **(C)**. n = 18 sections from 6 mice. (**E)** Tubulin polymerization by tau is compromised by HFD. Tubulin polymerization efficiency was measured by tau extracted from the heat stable fraction of the hippocampus. Values represent means ± SEM, n = 18 (3 technical replicates from 6 mice in each group). **(F)** HFD induces tau oligomerization. High molecular weight tau species (arrowheads, approximately 100, 140 and 250 kDa) were detected using the PS396 tau antibody (lower panel). **(G)** Quantification of high-molecular weight tau species as in **(F)**. n = 6 mice. (**H** and **I**) HFD reduces tau solubility. Tau in Tris-HCl-soluble (TS), Triton X100-soluble (TX), sarkosyl-soluble (SS), and sarkosyl-insoluble (SI) fractions were detected by TauC antibody (**H**) and quantified **(I)**. n = 6 mice. Graphs in **B, D, G, I, K** represent means ± SEM and *p* values are indicated in the graphs. Two-way ANOVA followed by Tukey-HSD (**B, D, G**) and Mann-Whitney U test (**I**). Scale bar: 500 μm (**C**).

GSK-3ß, which is one of key tau kinases associated with tauopathy, phosphorylates tau at Ser396 and Ser404 (Ser396/Ser404-phospho-tau) (Cavallini et al., 2013). Increased levels of Ser396/Ser404-phospho-tau were also observed in both Tg601 and NTg mice after HFD treatment (Figure 2A and B and Figure 2—figure supplement 1A). However, GSK-3ß phosphorylation at Tyr216 (positive regulation) and Ser9 (negative regulation) was both increased under HFD treatment. Considering increased Ser396/Ser404-phospho-tau, GSK-3β appeared to be activated on net regulation balance (Figure 2A and B). Other regulators of tau phosphorylation, including cyclin-dependent kinase 5, p38 mitogen-activated protein kinase, c-Jun N-terminal kinase and protein phosphatase 2A, did not show any changes after HFD treatment (Figure 2—figure supplement 2).

Tau hyperphosphorylation exerts tau pathologies, which include impaired tubulin polymerization and the formation of tau oligomers and sarkosyl-insoluble species (Di et al., 2016; Liu et al., 2007). Tubulin polymerization activity of tau in the heat stable fraction of the hippocampus was increased in NTg mice compared with Tg601 mice (Figure 2E). HFD treatment reduced the rate of tubulin polymerization in both Tg601 and NTg mice (Figure 2E). High-molecular weight multimeric or oligomeric phospho-tau, which is associated with tau pathology and synaptic loss (Kambe et al., 2011), appeared in HFD-treated NTg mice; this phenomenon was more obvious in HFD-treated Tg601 mice (Figure 2F and G). Moreover, sarkosyl-insoluble transgenic and endogenous tau was accumulated in HFD-treated Tg601 mice (Figure 2H and I).

The T2DM *db/db* genetic mouse model has displayed impaired hippocampal spatial learning and elevated glucocorticoid levels (Li et al., 2002; Livingstone et al., 2009; Stranahan et al., 2008). As expected, SGK1 upregulation and activation was observed in the hippocampi of *db/db* mice (Figure 2—figure supplement 3A, B). Consistent with increased SGK1 activation, increased Ser214-phospho-tau and increased GSK-3ß-associated Ser396/Ser404-phospho-tau were also observed, while total tau was unchanged (Figure 2—figure supplement 3A). The tubulin polymerization rate was reduced in *db/db* mice, corresponding to levels of tau phosphorylation (Figure 2—figure supplement 3C). In the hippocampi of mice treated with streptozotocin (STZ), which causes higher blood glucose levels and elevated glucocorticoid levels (Elahi et al., 2016a; Revsin et al., 2008), increased Ser214-phospho-tau and increased SGK1 phosphorylation at Thr256 and Ser422 was observed (Figure 2—figure supplement 4). Again, these results suggested that elevated glucocorticoids and hyperglycemia increasingly activate SGK1, leading to increased Ser214-phospho-tau levels.

**Figure 2-figure supplement 1.**
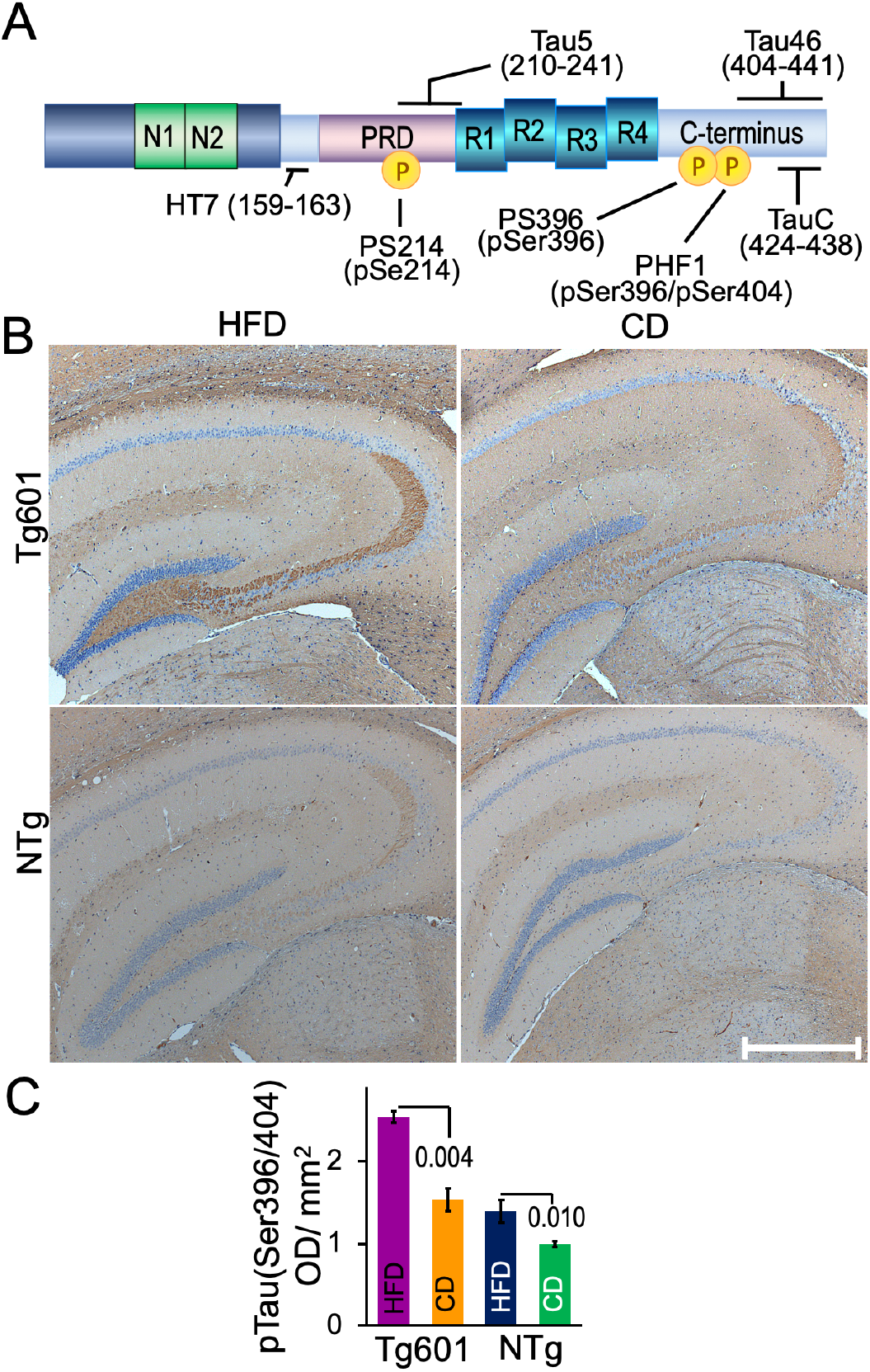
GSK-3ß-associated phosphorylation of tau is elevated by HFD. **(A)** Illustration of the human full-length 2N4R tau and epitopes of the anti-tau antibodies used in this study. The N- and C-terminal ends, two exons (N1, N2), proline-rich domain (PRD), and 4 repeats (R1-R4) are presented accordingly. **(B)** Immunohistochemistry of the hippocampus with the PHF1 antibody. **(C)** Quantification of PHF1 immunoreactivity. Graph is represented as the means ± SEM (n = 18 sections from 6 mice in each group) and were analyzed by two-way ANOVA followed by Tukey HSD **(C)**. *p*-values are indicated in the graphs when significantly different. Scale bar: 500 μm (**B**).

**Figure 2-figure supplement 2.**
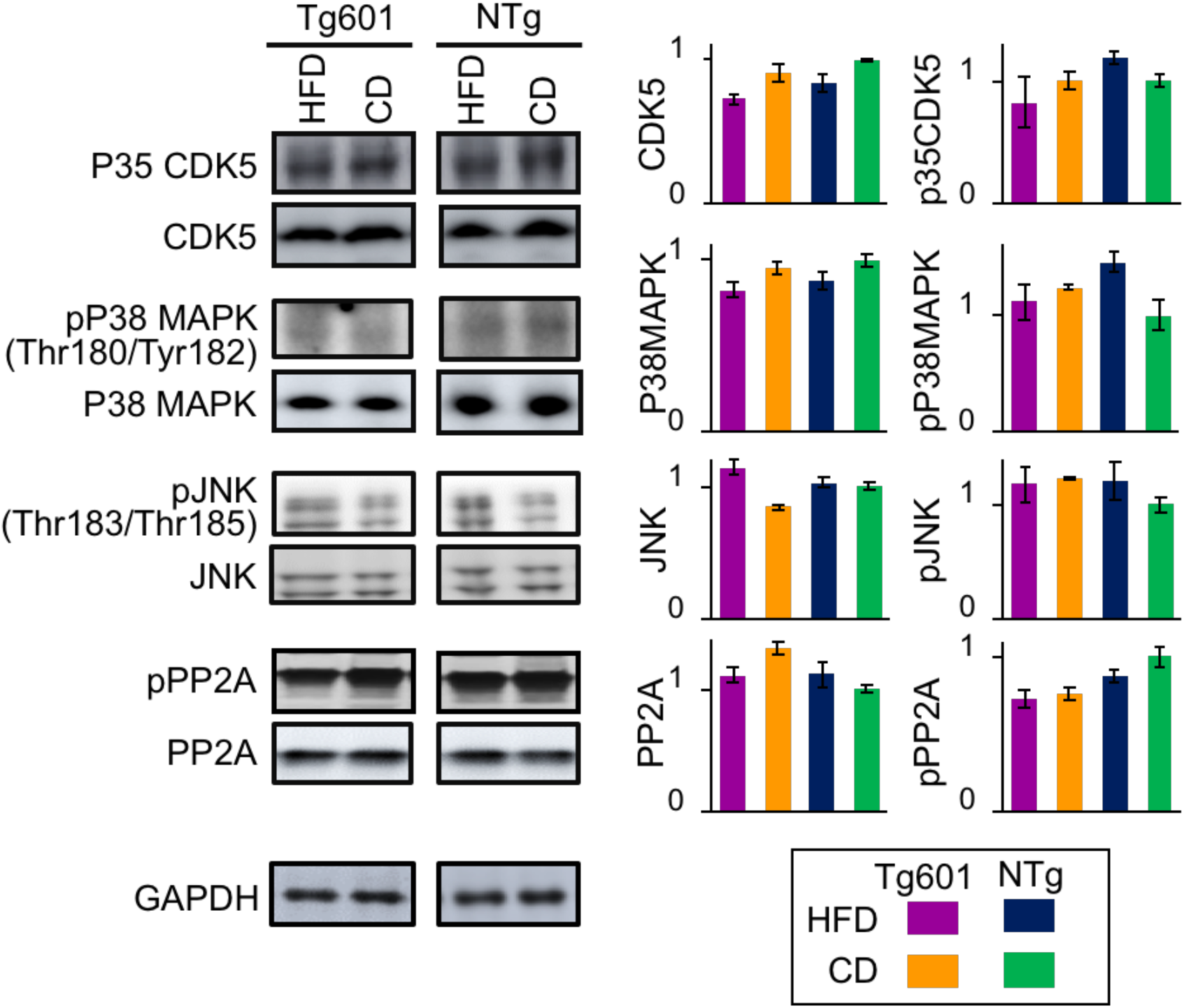
Expression of tau kinases and phosphatases in the hippocampus of HFD-treated mice. Effect of HFD on hippocampal tau kinases including CDK5, p38 MAPK, c-Jun N-terminal kinase (JNK) and protein phosphatase 2A (PP2A) in the hippocampi of Tg601 and NTg mice. Western blotting analysis was performed with the indicated antibodies. Quantification of different tau kinases and PP2A. Graphs are represented as the means ± SEM (n = 6 mice in each group) and were analyzed by Mann-Whitney U test. All data were not significantly different.

**Figure 2-figure supplement 3.**
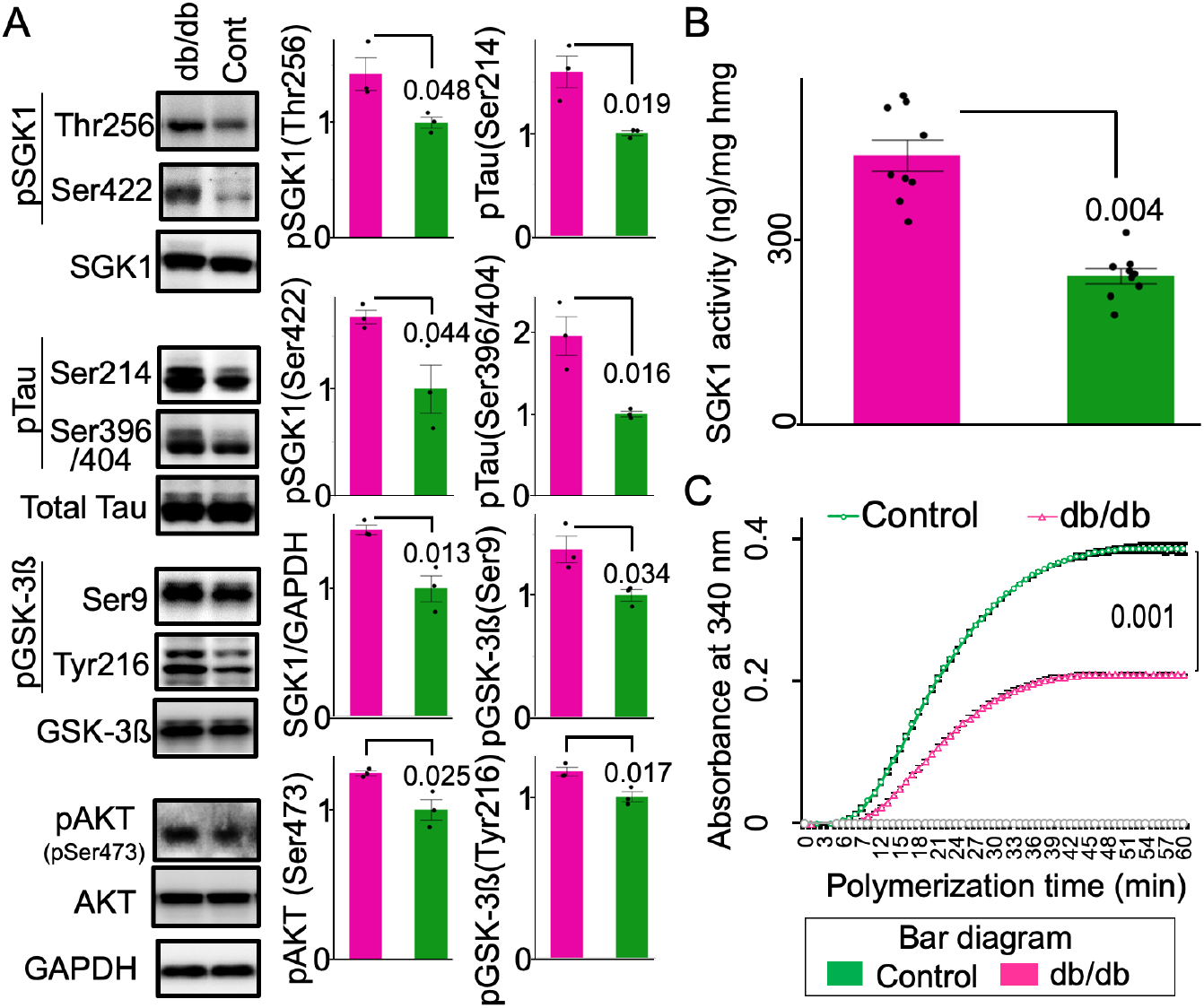
SGK1 kinase activity is elevated, and tubulin polymerization activity is reduced in the *db/db* mouse brain. **(A)** Phosphorylation of SGK1 and tau is increased in the *db/db* hippocampus. **Densitometric** quantification of the indicated proteins. n = 3 mice. **(B)** SGK1 kinase activity in the *db/db* mouse brain. An ADP-Glo kinase assay was performed using immunopurified SGK1 and SGKtide. n = 3 mice in each group and 3 replicates for each mouse. **(C)** Tubulin polymerization efficiency of tau extracted from the heat-stable fraction of *db/db* and control brains. n = 3 biological replicates. Graphs are represented as the means ± SEM. A paired Student’s *t*-test.

**Figure 2-figure supplement 4.**
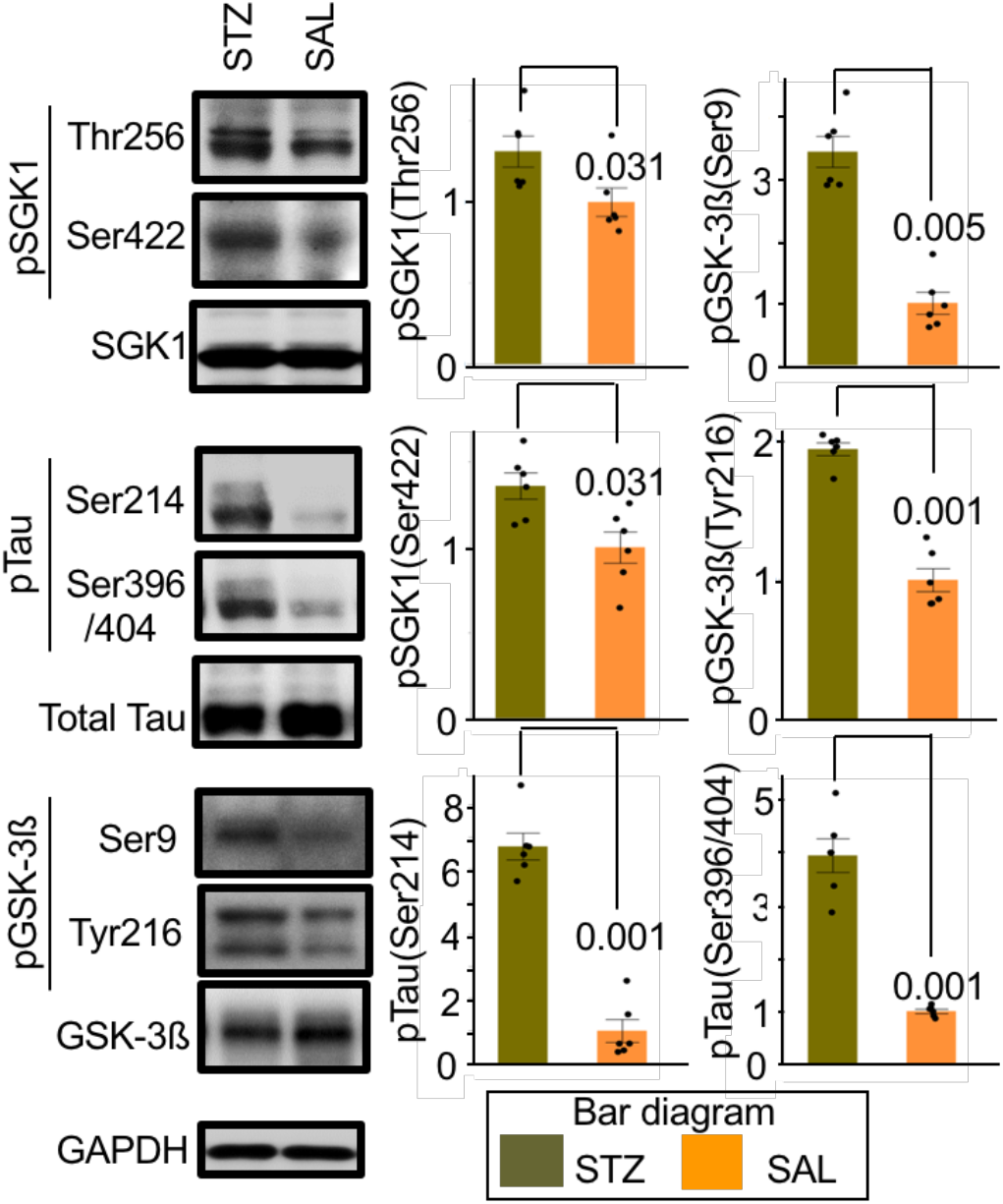
STZ administration stimulates SGK1 activity and tau phosphorylation in the mouse hippocampus. Western blot analysis of STZ- or saline (SAL)-injected mouse hippocampus using the indicated antibodies. Graphs represent quantification of immunoreactivity. Graphs are represented as the means ± SEM (n = 6 mice in each group). Mann-Whitney U test and paired Student’s *t*-test based on distribution analysis.

Elevated SGK1 expression by recombinant adeno-associated virus (AAV) infection in the hippocampi of Tg601 mice caused increased phosphorylation of Ser214-phospho-tau and SGK1 phosphorylation at Thr256/Ser422 as well as Ser396/Ser404-phospho-tau and GSK-3ß phosphorylation at Tyr216/Ser9 (Figure 3A-C). TUNEL-positive neurons were also detected in exogenous SGK1-positive regions (Figure 3D). Consistent with the increased pathogenesis-associated tau phosphorylation, exogenous SGK1 expression in Tg601 mice impaired the spatial memory/learning and recognition, but not affected anxiety like behavior (Figure 3E-H).

**Figure 3.**
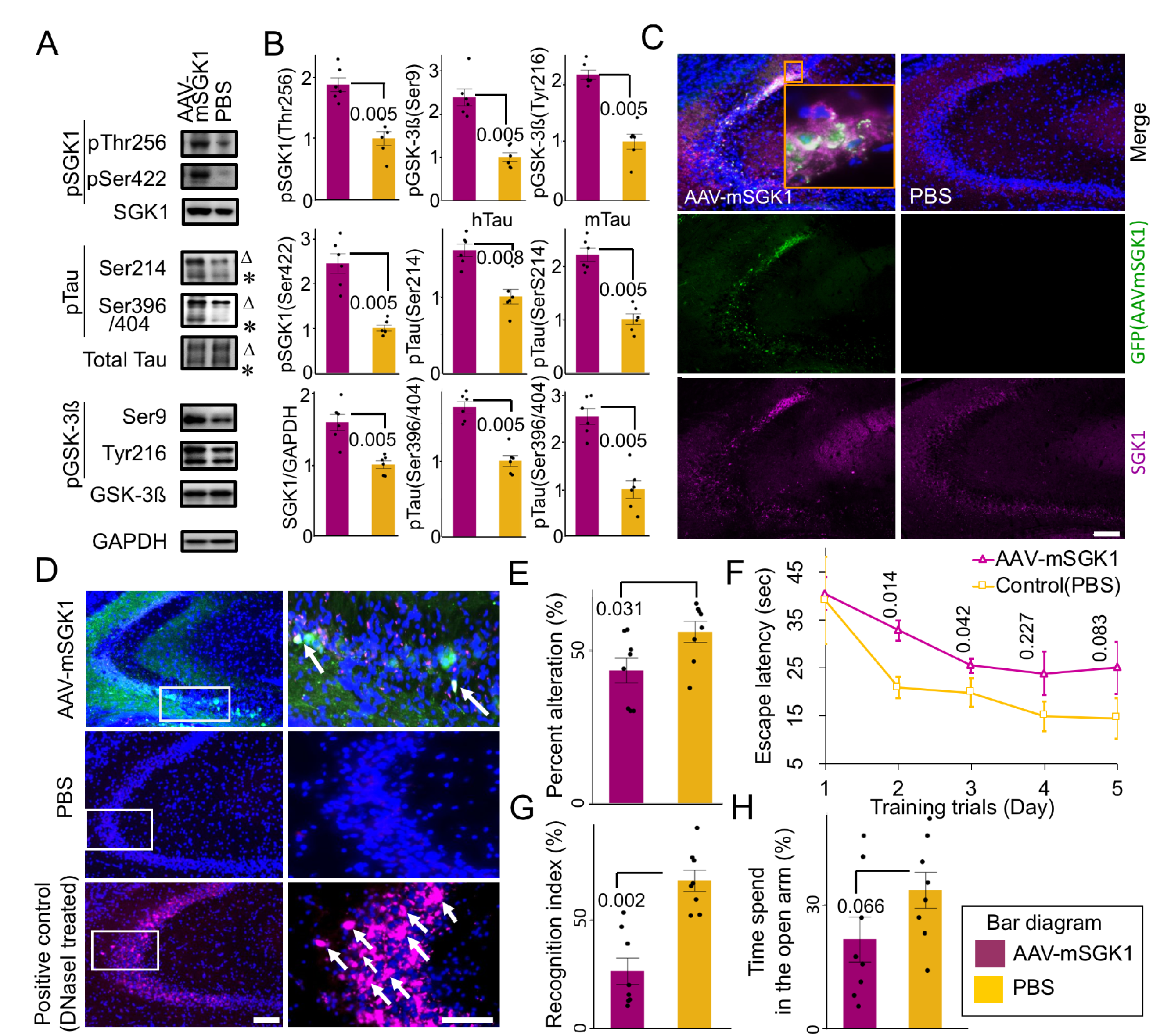
Exogenous SGK1 expression in the hippocampal neurons leads to neurodegeneration and impaired cognitive functions in Tg601 mice. **(A, B)** Representative results of western blotting **(A)** and densitometric quantification **(B)** of tau and GSK-3ß phosphorylation (n = 6 mice, arbitrary units) in Tg601 mouse hippocampus injected with mock PBS or AAV-mSGK1-2A-EGFP. Triangles and asterisks indicate human Tau (hTau) and mouse Tau (mTau), respectively. **(C)** SGK1 immunofluorescence (red) in Tg601 mouse hippocampal CA2/CA3 regions, where AAV-mSGK1-2A-EGFP or PBS was injected. EGFP (green) dissociated from SGK1 by the self-cleavage of a 2A peptide marked the SGK1-overexpressing regions. Scale bar: 100 μm. **(D)** TUNEL assay in the hippocampal sections, treated as in (**A-C**). DNase I-treated section served as a positive control. TUNEL/EGFP-positive-cells were indicated by arrows. Scale bars: 100 μm (left) and 50 μm (right). **(E-H)** Elevated SGK1 impairs learning and memory. Twenty-one days after Tg601 hippocampal injection of the AAV-mSGK1-2A-EGFP, cognitive performance was investigated (n = 8 mice in each group). **(E)** mSGK1 overexpression worsens spatial memory. SGK1 overexpression reduced spontaneous alternation behavior of Tg601 mice in the Y maze test. **(F)** SGK1 overexpression worsens spatial learning. Hidden platform trials were performed in the Morris water maze. AAV-mSGK1-injected mice had longer escape latencies than PBS-injected mice in day 2 and 3. **(G)** SGK1 overexpression reduces recognition of novel objects. Recognition index (%) were expressed as means ± SEM (3 trials with 8 mice in each group, see Materials and Methods for detail). **(H)** Anxiety test. Fear feelings were assessed by the time spent in open arms in an elevated plus-maze test. AAV-mSGK1-injected mice showed no significant changes in this test.

### Diabetes-associated dexamethasone and high blood glucose activate SGK1

As a single factor is unlikely to be involved in the hippocampal SGK1 activation in the HFD-treated mice, we set up three conditions associated with T2DM, which included conditions wherein cells were exposed to dexamethasone (DXM, a glucocorticoid analog), high glucose (a hyperglycemia-like condition) and palmitic acid conjugated with bovine serum albumin (PA-BSA, a hyperlipidemia-like condition). DXM upregulated and activated SGK1 in human neuroblastoma SH-SY5Y cells expressing transgenic tau, leading to increased Ser214-phospho-tau in both a dose- and time-dependent manner (Figure 4A and B and Figure 4—figure supplement 1). In addition, DXM treatment increased GSK-3ß expression and activation by increasing Tyr216 phosphorylation (Figure 4—figure supplement 1A and B). Consistent with increased SGK1 and GSK-3ß activation, the tubulin polymerization activity of tau taken from DXM-treated cells was decreased (Figure 4C). Moreover, high-molecular weight Ser396-phospho-tau was elevated in DXM-treated cells (Figure 4D). High glucose treatment also increased SGK1 and GSK-3ß activation and reduced the tubulin polymerization activity of tau, although these effects were milder than what was observed with DXM treatment (Figure 4A-C and Figure 4—figure supplement 1A and B). In contrast, palmitic acid (PA) treatment, which mimics fatty acid-mediated insulin resistance, did not affect SGK1 expression or activation. However, PA treatment resulted in a marked reduction in levels of phosphorylated AKT at Ser473 (pSer437 AKT) and phosphorylated GSK-3ß at Ser9 (pSer9 GSK3ß), suggesting that reduced AKT activation results in increased GSK-3ß activation and subsequent tau phosphorylation apart from SGK1 (Figure 4A and B and Figure 4—figure supplement 1A and B).

**Figure 4.**
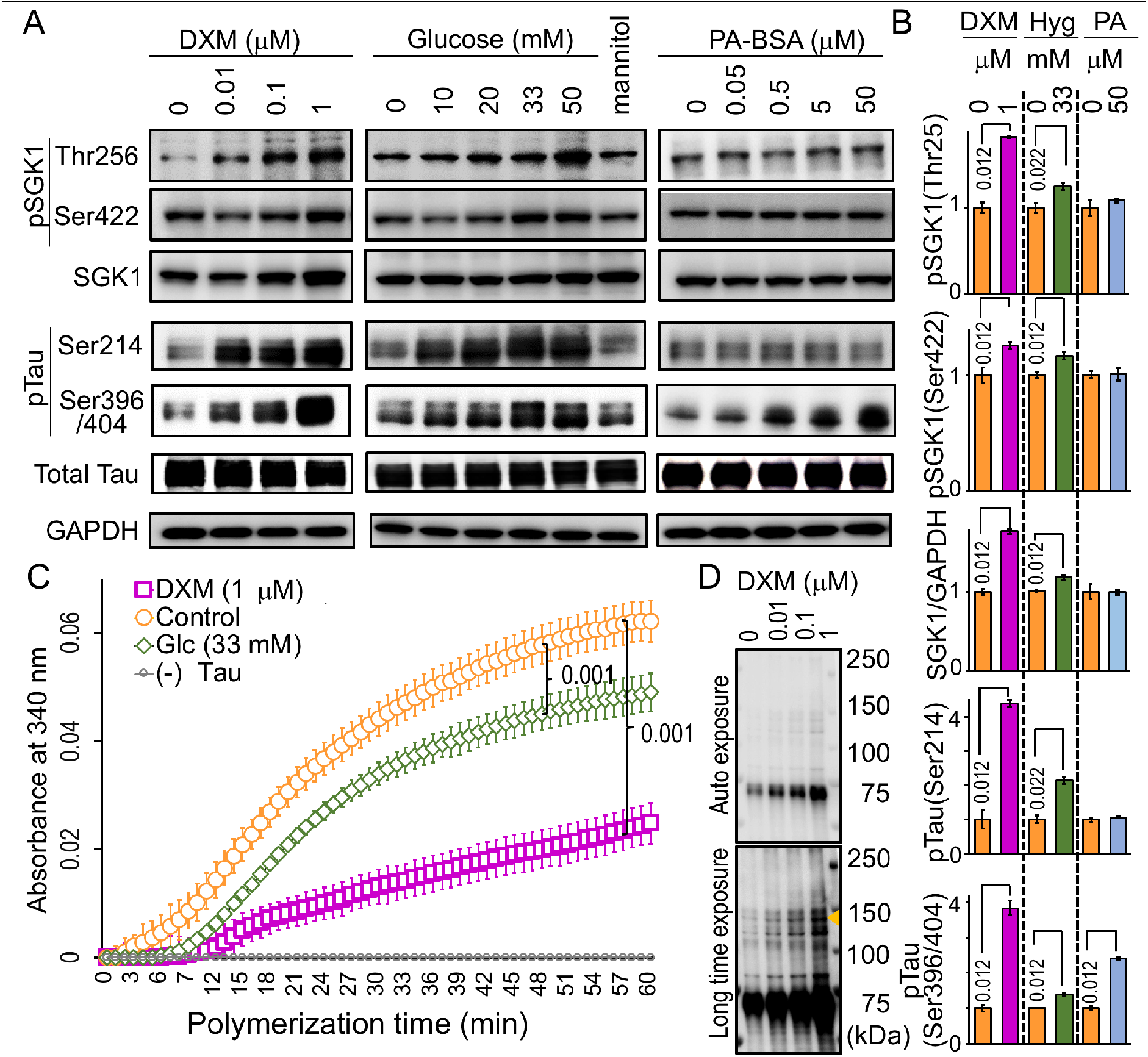
DXM and glucose, but not fatty acids, activate SGK1 in cultured cells. **(A)** DXM and high glucose stimulate SGK1 activation and tau phosphorylation in SH-SY5Y cells stably expressing tau. Cells were treated with increasing concentrations of DXM for 3 hrs, glucose for 6 hrs and PA-BSA for 48 hrs to mimic various consequences of HFD. Mannitol was used as an osmolarity control for glucose. **(B)** Quantification of the indicated protein levels treated with or without 1 μM DXM, 33 mM glucose, and 50 μM PA-BSA as in **A**. n = 5 biological replicates. **(C)** Tau from cells treated with DXM or high glucose impairs tubulin polymerization. n = 5 biological replicates. DXM-non-treated samples were used as a control. **(D)** DXM promotes high molecular weight tau formation (arrowhead). Blots were detected with PS396 antibody. Graphs in **B** represent means ± SEM and *p* values are indicated in the graphs. Mann-Whitney U test **(B, C)**.

**Figure 4-figure supplement 1.**
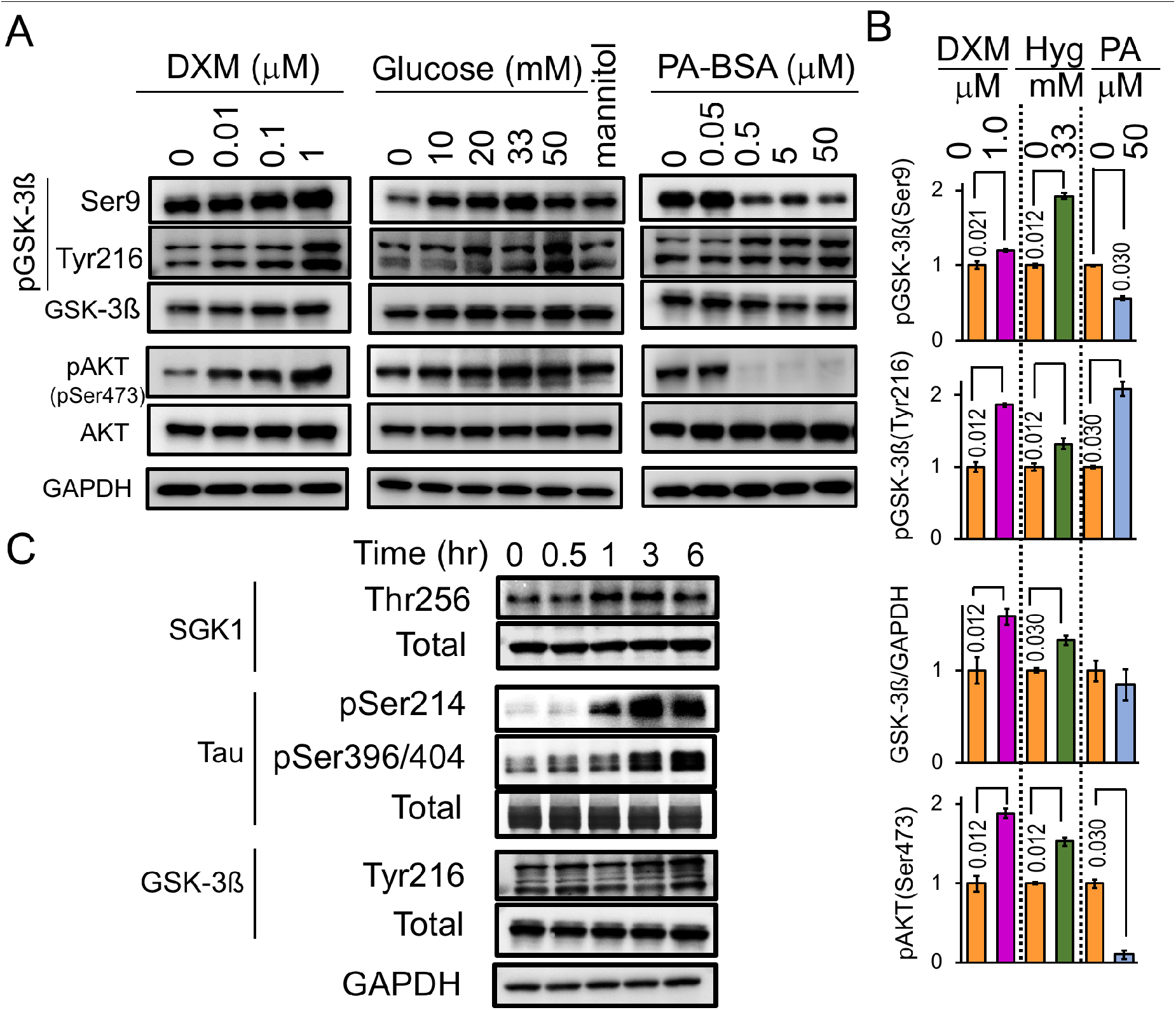
Effect of DXM, high glucose and palmitic acid on GSK-3ß and AKT. **(A, B)** Dose-dependent effects of DXM, high glucose, and PA-BSA on GSK-3ß and AKT in SH-SY5Y cells. Western blot data **(A)** and quantification analysis **(B)** of phosphorylated GSK-3ß and AKT with the indicated treatments. **(C)** Time-dependent effects of DXM on SGK1 and GSK-3ß activation and tau phosphorylation. SH-SY5Y cells were treated with 1 μM DXM for the indicated time periods. Graphs are represented as the means ± SEM (n = 5 biological replicates in each group). Mann-Whitney U test.

### SGK1-mediated tau phosphorylation at Ser214 and GSK-3ß activation

To determine the possibility that SGK1 activates GSK-3ß, we next examined the effects of SGK1 inhibition on GSK-3ß activation and tau phosphorylation. Knockdown of SGK1 in SH-SY5Y cells stably expressing tau reduced levels of phosphorylated GSK-3ß at Tyr216 (pTyr216 GSK3ß) (Figure 5A and B), a key phosphorylation site that increases GSK-3ß activity, leading to a reduction in Ser214-phospho-tau levels caused by SGK1 activity and Ser396/404-phospho-tau levels caused by GSK-3ß activity (Figure 5A and B). In contrast, levels of pSer9 GSK-3ß and pSer473 AKT were unaffected by SGK1 knockdown, suggesting that GSK-3ß pSer9 phosphorylation is predominantly regulated by AKT or other kinases (Figure 5A and B).

**Figure 5.**
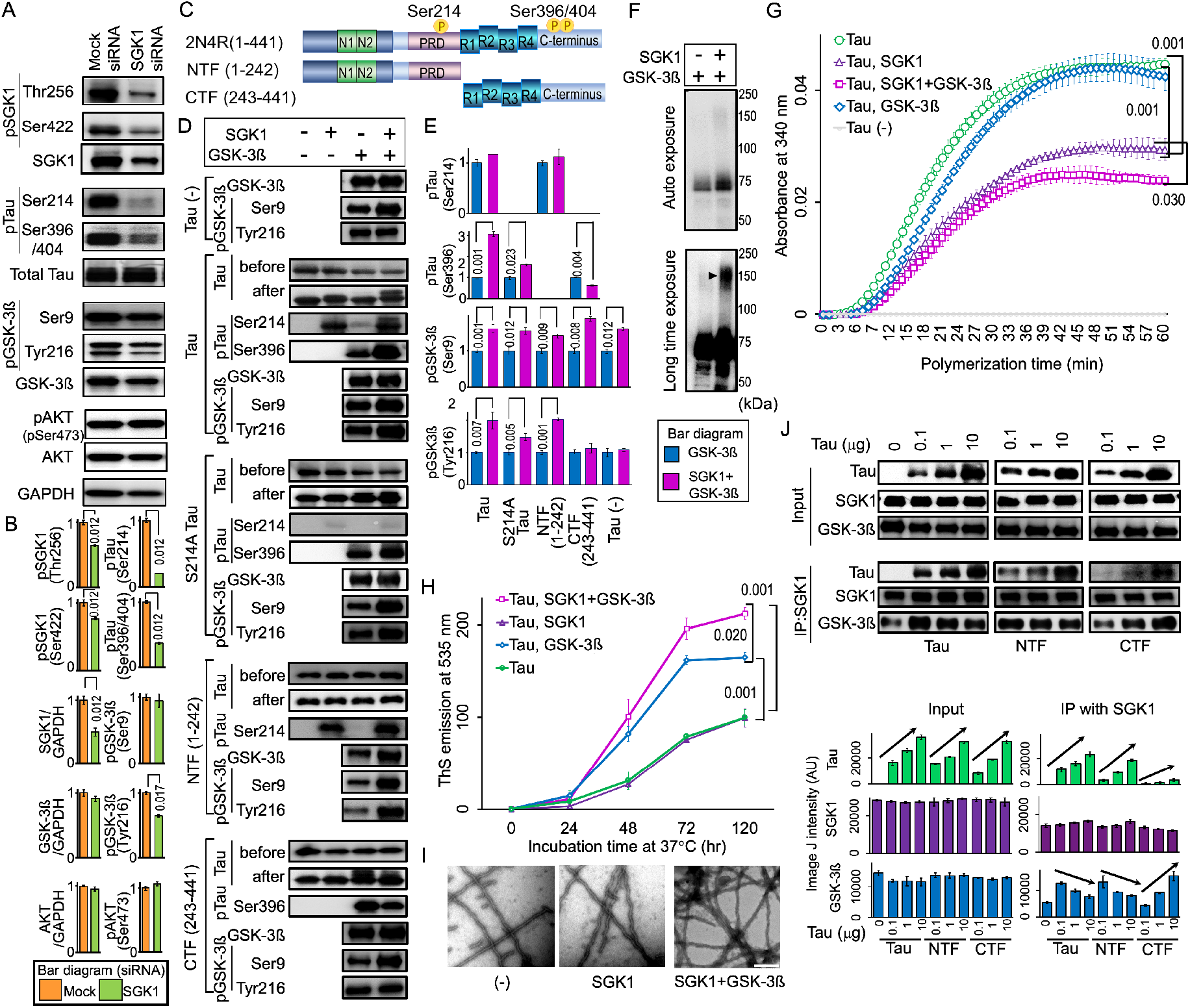
SGK1 activates GSK-3ß, leading to GSK-3ß-associated tau phosphorylation where tau functions as a scaffold for SGK1-mediated GSK-3ß activation. **(A)** SGK1 suppression downregulates GSK-3ß phosphorylation at pTyr216. The indicated proteins were analyzed using SH-SY5Y cells stably expressing tau. (**B)** Quantification of the indicated protein levels. Graphs represent means ± SEM and *p* values are indicated in the graphs (n = 5 biological replicates, Mann-Whitney U test). **(C-E)** The N-terminus of tau is required for GSK-3ß phosphorylation at pTyr216. Kinase assays were performed using recombinant SGK1 and/or GSK-3ß along with wild-type tau, Ser214Ala tau, N-terminal (NTF, 1-242 aa) or C-terminal (CTF, 243-441 aa) tau fragments that were used as substrates. **(C)** A schematic representation of the tau variants used in the kinase assay. **(D)** tau blots before and after kinase reactions. **(E)** Quantification of the indicated protein levels (n = 4 biological replicates). **(F)** SGK1 promotes high molecular weight multimeric tau formation by GSK-3ß. Western blots were performed after kinase reactions as in **(D)** using the PS396 antibody. **(G)** GSK-3ß exacerbates SGK1-mediated suppression of microtubule polymerization activity of tau. n = 4 biological replicates. **(H)** SGK1 promotes tau aggregation and fibril formation by increasing GSK-3ß activity. tau was prepared from purified recombinant tau protein treated with or without SGK1 and/or GSK-3ß. TAU (15 μM) and heparin (0.1 mg/ml) were incubated at 37°C for the indicated periods of time using a ThS assay. n = 5 biological replicates. **(I)** Transmission electron micrograph images of the tau fibrils treated as in **(H). (J)** SGK1 directly binds to GSK-3ß and indirectly binds to tau’s CTF via GSK-3ß. Co-immunoprecipitation of SGK1 and GSK-3ß using an anti-SGK1 antibody after the kinase reaction as in **(C)**. The kinase reaction was performed in the presence of increasing amounts of tau or its fragments. Input was 10% of the volume used for IP. Graphs represent quantification of the indicated proteins. n = 3 biological replicates. IP, immunoprecipitation. Data ± SEM and *p* values are indicated in the graphs. Mann-Whitney U test (**E, G**) and two-way ANOVA followed by Tukey-HSD Graphs in **G** represent means (**H**). Scale bar: 200 nm (**I**).

SGK1 is shown to directly phosphorylate tau at Ser214 (Figure 5C-E, Figure 5—figure supplement 1 and Figure 5—figure supplement 2A). In addition, SGK1 augmented the kinase activity of GSK-3ß toward increasing tau phosphorylation (Figure 5C-E and Figure 5—figure supplement 2B) and GSK-3ß-mediated tau dimerization (Figure 5F). As previous reports suggest, the tubulin polymerization activity of tau after the SGK1 kinase reaction indicated that tau phosphorylated at Ser214 does not readily interact with tubulin (Virdee et al., 2007; Yang et al., 2006), which was further enhanced by GSK-3ß treatment (Figure 5G). While SGK1 had no effect on heparin-induced tau aggregation, the combination of SGK1 and GSK-3ß resulted in increased tau aggregation compared with the activity of GSK-3ß alone (Figure 5H). The ultrastructural observation of resultant tau fibrils in a Thioflavin-S (ThS) assay confirmed that there was increased fibril formation when tau was treated with both SGK1 and GSK-3ß (Figure 5I).

A putative SGK1 phosphorylation motif is present at Ser9 of GSK-3ß, which may negatively regulate its kinase activity (Krishnankutty et al., 2017; Murray et al., 2005). In *in vitro*, SGK1 increased pSer9 GSK-3ß phosphorylation in the absence of tau, whereas activation-associated pTyr216 GSK-3ß was unaffected, which ultimately leads to a reduction of GSK-3ß activity (tau (-) in Figure 5D and E). A similar result was also observed when the C-terminal tau fragment (CTF, 243-441 aa) (Matsumoto et al., 2015) lacking the SGK1 phosphorylation motif was used in a kinase reaction (Figure 5D and E). However, in the presence of the N-terminal tau fragment (NTF, aa 1-242) or full-length tau, the phosphorylation of GSK-3ß both at Tyr216 and Ser9 was enhanced, and ultimately lead to activated GSK-3ß kinase (Figure 5D and E). These results suggest that the NTF is required for SGK1-mediated GSK-3ß Tyr216 phosphorylation.

SGK1-mediated GSK-3ß activation was also observed with tau containing a Ser214Ala mutation (Figure 5D and E), which ultimately increased tau phosphorylation at Ser396. The increase in tau phosphorylation at Ser396 was also supported by phos-tag Western blot analysis (Figure 5—figure supplement 2C and D), which also revealed that SGK1 phosphorylates tau at an additional site other than Ser214 (Figure 5—figure supplement 2C and Figure 5—figure supplement 3). However, only the phosphorylation of tau at Ser214 weakened the tubulin polymerization activity of tau (Figure 5—figure supplement 2E). SGK1 did not form a complex with tau during the phosphorylation reaction as is observed with GSK-3ß (Figure 5—figure supplement 2F). In contrast, SGK1 was shown to bind to GSK-3ß, which facilitated the association of SGK1 with tau through the NTF region (Figure 5J and Figure 5—figure supplement 2F). GSK-3ß directly bound to the CTF (Figure 5—figure supplement 2F). These results suggest that SGK1 activates GSK-3ß by forming a scaffold-like complex with tau and GSK-3ß, which in turn stimulates tau aggregation, likely through enhanced phosphorylation of tau at GSK-3ß-responsible sites (Figure 5—figure supplement 3).

**Figure 5-figure supplement 1.**
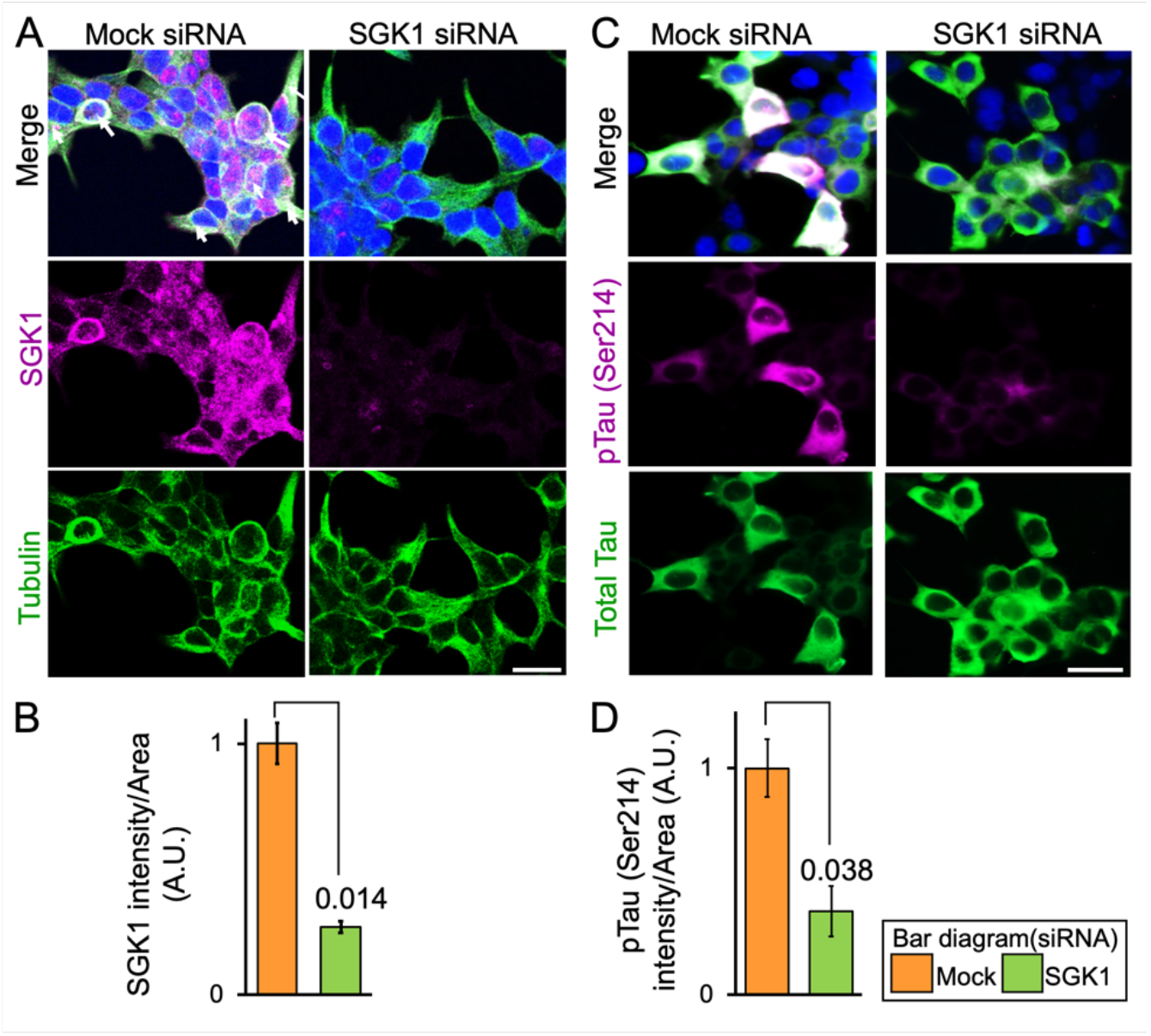
Knockdown of SGK1 decreases tau phosphorylation at Ser214. **(A, C)** Confocal immunofluorescence microscopic images of SH-SY5Y cells stably expressing tau treated with SGK1 siRNA or control siRNA. Bars, 50 μm. **(B, D)** Graphs represent quantification of immunofluorescent intensity in the tubulin- or tau-positive areas. Scale bar: 50 μm (**A, C**). Graphs are represented as the means ± SEM (n = 6 images from 5 independent samples in each group). A paired student *t*-test.

**Figure 5-figure supplement 2.**
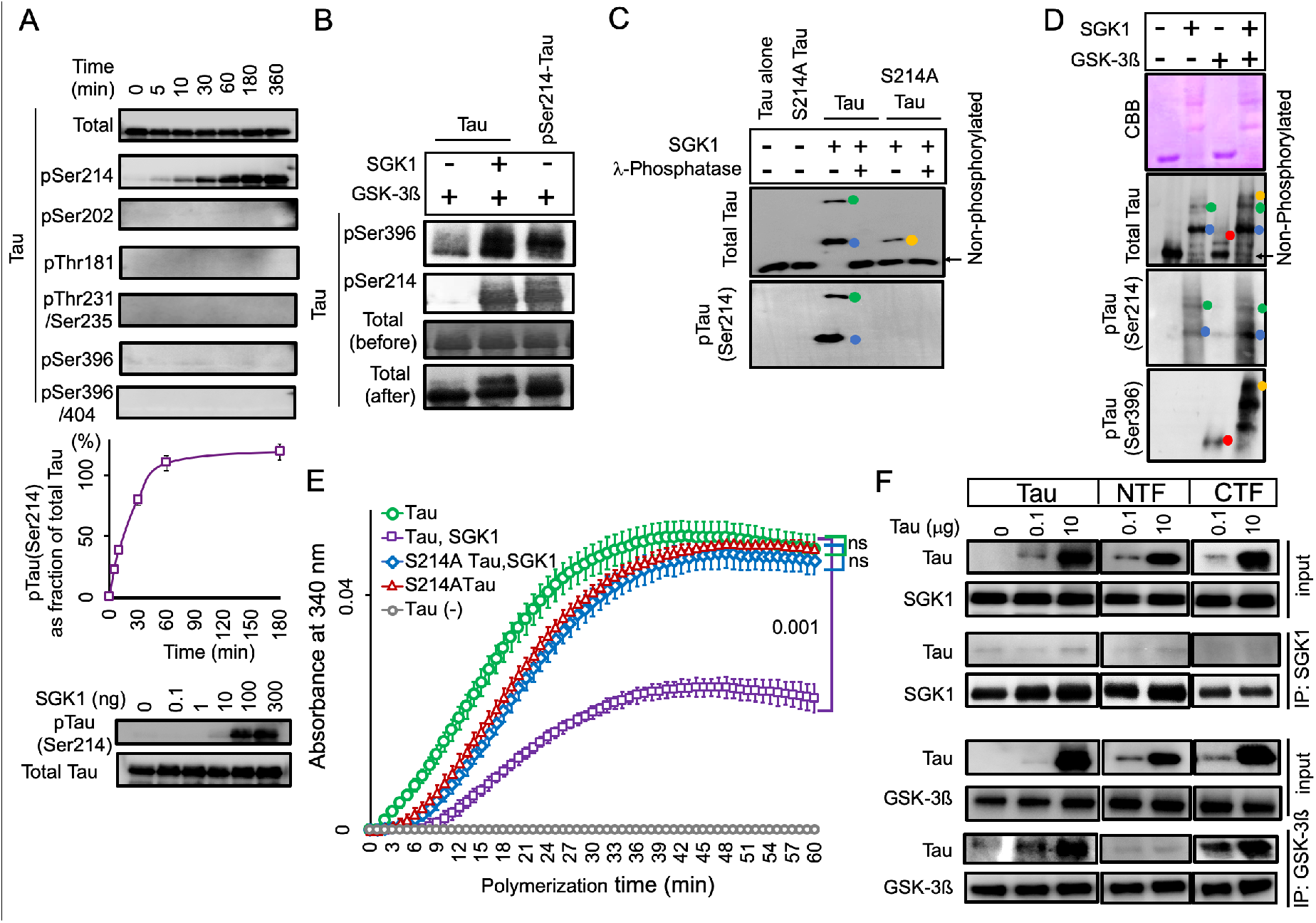
SGK1 directly phosphorylates tau at Ser214. **(A)** SGK1 phosphorylates tau at Ser214 *in vitro*. Recombinant 2N4R tau treated with SGK1 for the indicated periods of time were analyzed by Western blot with the indicated phospho-tau-antibodies. Total tau signals were confirmed by an anti-tau antibody (tau5). The Ser214 phosphorylation reaction reaches a plateau at 60 min (middle). n = 2 technical replicates. Dosedependent kinase activity of SGK1 on Tau using different doses of SGK1 in a 1-hr kinase reaction (low). **(B)** pSer214 has a priming effect on the phosphorylation of pSer396 by GSK-3ß. For pSer214-tau preparation, tau pretreated with (lane 3) or without SGK1 (lanes 1 and 2) was heat-inactivated and immunoprecipitated with an anti-tau antibody. Precipitated tau was further incubated with GSK-3ß (lanes 1 and 3) or GSK-3ß and SGK1 (lane 2). Tau phosphorylation at Ser396 was enhanced by pre-existing pSer214 (lane 1 *vs.* lane 3). **(C)** Phostag Western blot analysis reveals that SGK1 phosphorylates an uncharacterized site (X) along with Ser214 in tau. Recombinant wild-type or Ser214Ala mutated tau was incubated with SGK1 in the presence or absence of λ phosphatase. Colored dots indicate tau with phosphorylation at Ser214 (blue), tau with phosphorylation at two sites (Ser214 and X, green), and tau with phosphorylation at X (orange). **(D)** Phos-tag Western blot analysis were performed after the kinase reaction using SGK1 and/or GSK-3ß along with full-length tau. Colored dots indicate tau with phosphorylation at Ser214 by SGK1 (blue), tau with phosphorylation at two sites (Ser214 and X) by SGK1(green), tau with phosphorylation at Ser396 by GSK-3ß (red), and tau that exhibited hyperphosphorylation at sites affected by both SGK1 and GSK-3ß (orange). **(E)** Analysis of the effect of SGK1-mediated Ser214 phosphorylation on suppression of microtubule polymerization activity of tau. **(F)** SGK1 does not bind to tau while GSK-3ß binds to tau CTF. SGK1 or GSK-3ß was immunoprecipitated in the presence or absence of full-length or truncated tau using anti-SGK1 or anti-GSK-3β antibody. Co-precipitated tau is detected with anti-tau antibodies (Tau5 and Tau46). Graphs are represented as the means ± SEM (n = 4 biological replicates). Mann-Whiney U test (**E**).

**Figure 5-figure supplement 3.**
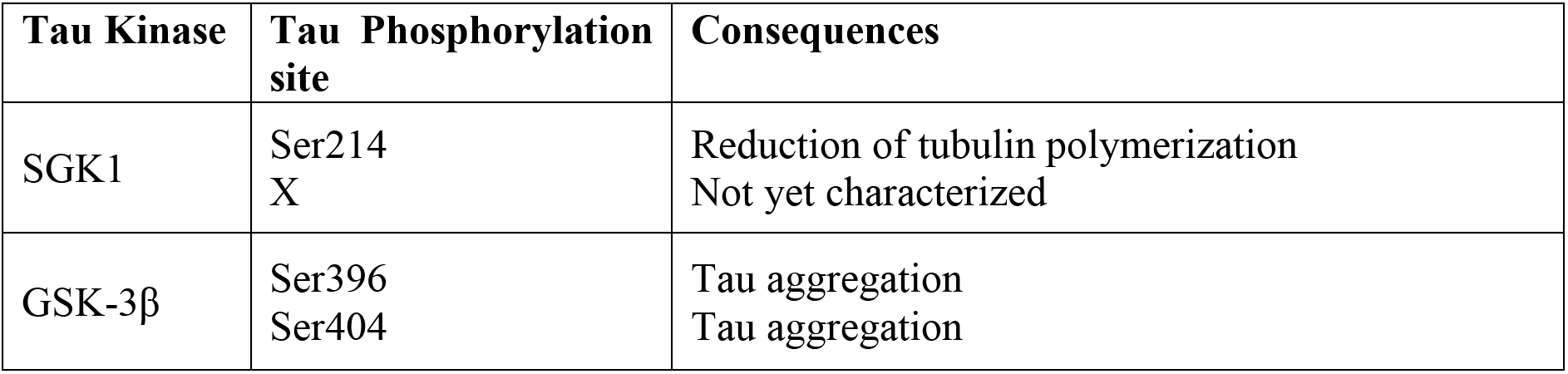
Tau kinase, phosphorylation sites, and consequences by phosphorylation.

### EMD638683 suppresses GSK-3ß activation and tau phosphorylation by inhibiting SGK1

EMD638683, a potent inhibitor of SGK1, was previously reported to have anti-hyperglycemic and anti-hypertensive activity (Ackermann et al., 2011; Gan et al., 2018; Li et al., 2016). *In vitro* kinase assays using recombinant SGK1, GSK-3ß and tau proteins showed a dosedependent inhibition of Ser214-phospho-tau (Figure 6A-C), but not Ser396-phospho-tau (Figure 6—figure supplement 1A), that was caused by EMD638683. Although EMD638683 did not directly affect GSK-3ß activity (Figure 6—figure supplement 1A), it reduced pTyr216 GSK-3ß levels and GSK-3ß-associated Ser396-phospho-tau levels in combination with SGK1 (Figure 6A-C). Moreover, EMD638683 reversed SGK1-mediated inhibition of the tau tubulin polymerization activity (Figure 6D).

**Figure 6.**
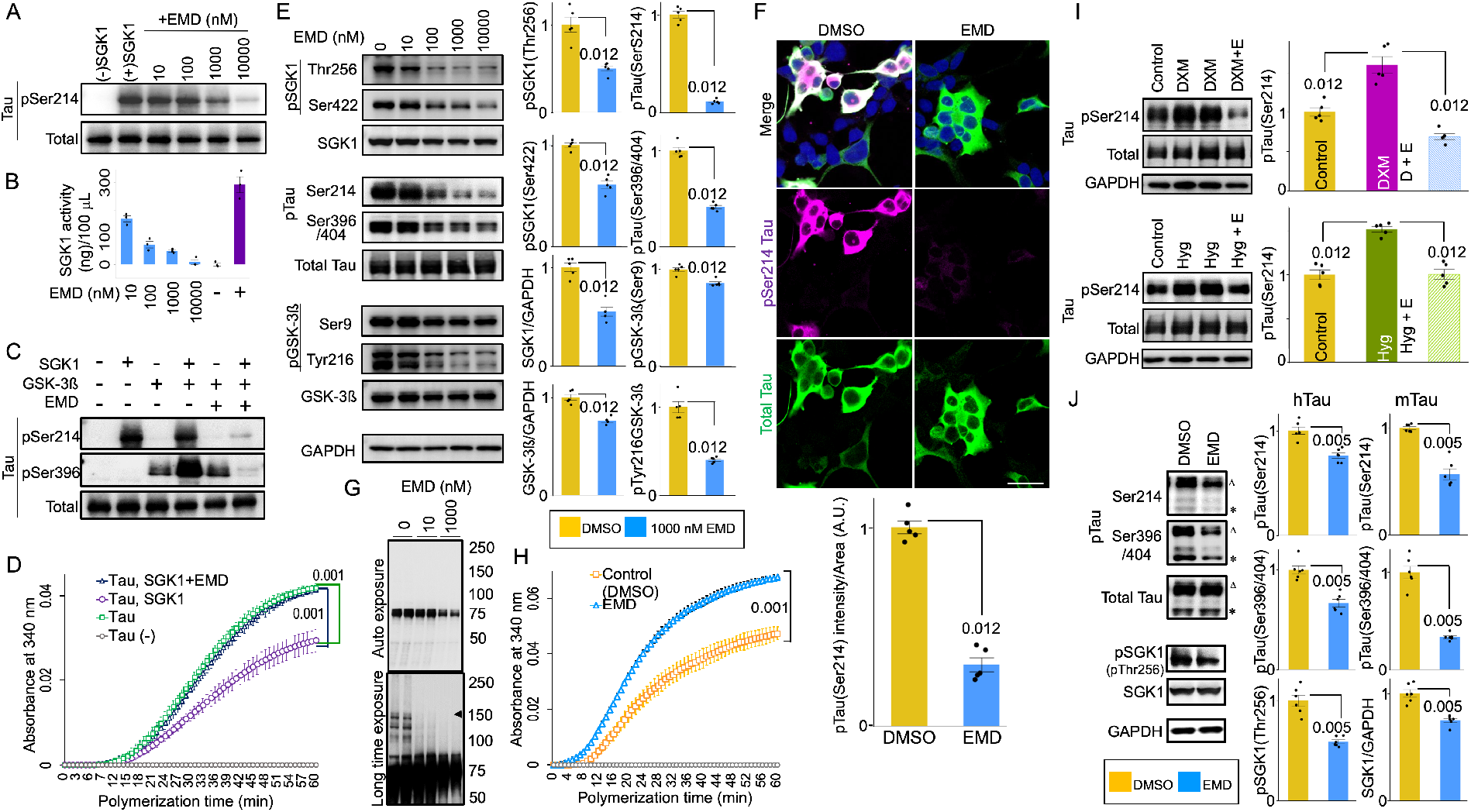
SGK1 inhibitor EMD638683 ameliorates GSK-3ß-mediated tau phosphorylation. **(A, B)** Dose-dependent inhibition of SGK1 by EMD638683. Phosphorylation of tau at Ser214 by SGK1 was monitored using recombinant SGK1 and recombinant tau **(A)**. SGK1 activity was monitored using the ADP-Glo kinase assay using recombinant tau as a substrate. n = 3 biological replicates **(B)**. **(C)** EMD638683 inhibits GSK-3ß kinase activity through SGK1. GSK-3ß kinase activity toward tau was evaluated using the pSer396 tau antibody. **(D)** EMD638683 restores the tubulin polymerization ability of tau in the presence of SGK1. n = 4 biological replicates. **(E)** EMD638683 inhibits both SGK1- and GSK-3ß-associated phosphorylation sites of tau. SH-SY5Y cells stably expressing tau were treated with the indicated concentrations of EMD638683 for 24 hrs. Graphs represent quantification of the indicated proteins. n = 5 biological replicates. **(F)** EMD638683 results in reduced Ser214-phospho-tau in SH-SY5Y cells stably expressing tau. Cells were treated with EMD638683 for 24 hrs. Graph represents quantification of pSer214 fluorescent intensities in tau-positive areas. n = 8 images from 5 biological replicates. **(G)** EMD638683 suppresses high molecular weight tau formation (arrowhead). Blots were detected using the PS396 antibody. **(H)** EMD638683 improves the tubulin polymerization ability of tau. n= 5 biological replicates. **(I)** DXM (upper panel)- or high glucose (lower panel)-mediated pSer214 tau phosphorylation is inhibited by EMD638683. SH-SY5Y cells stably expressing tau were treated with 1 μM DXM or 33 mM glucose for 6 hr in the presence (D + E or Hyg + E) or absence of 1 μM EMD638683. Graphs represent quantification of the indicated proteins. n = 5 biological replicates. **(J)** EMD638683 inhibits both SGK1- and GSK-3ß-associated phosphorylation sites of tau in the hippocampi of Tg601 mice. Tg601 mice were injected stereotactically with 5.0 mg/kg of EMD638683 in the hippocampi and sacrificed after 72hrs. Graphs represent quantification of the indicated proteins. n = 6 mice. EMD, EMD638683. Graphs in **B, D-F, H-J** represent means ± SEM and *p* values are indicated in the graphs. Mann-Whitney U test **(B, D, E, H, I)** and a paired Student *t*-test **(F, J)**. Scale bar: 50 μm **(F)**.

EMD638683 also inhibited SGK1 in SH-SY5Y cells stably expressing tau in dose- and time-dependent manners without affecting tau levels, cell survival or cytotoxicity (Figure 6E and F and Figure 6—figure supplement 1B-D). The EC_50_ of EMD638683 on pSer214 tau in SH-SY5Y cells was 99.3 nM (Figure 6—figure supplement 1E). EMD638683 treatment at 100 nM sufficiently suppressed the formation of high-molecular weight tau in SH-SY5Y cells and improved tau’s tubulin polymerization ability (Figure 6G and H). Moreover, EMD638683 significantly reduced pTyr216 GSK-3ß and Ser396/404-phospho-tau levels, while expression levels of tau were unaffected (Figure 6E). EMD638683 also suppressed Ser214-phospho-tau levels in cells treated with DXM or high glucose (Figure 6I). A reduced level of pSer214 and Ser396/404-phospho-tau was observed in the Tg601 mouse hippocampi after injection with EMD638683 (Figure 6J). Finally, the elevated levels of pSer214 tau and pSer256/pSer422 SGK1, forming the SGK1-GSK-3ß-tau tertiary complex, were observed in AD patients, strongly suggesting that SGK1-mediated tau phosphorylation is involved in the etiology of AD (Figure 7).

**Figure 6-figure supplement 1.**
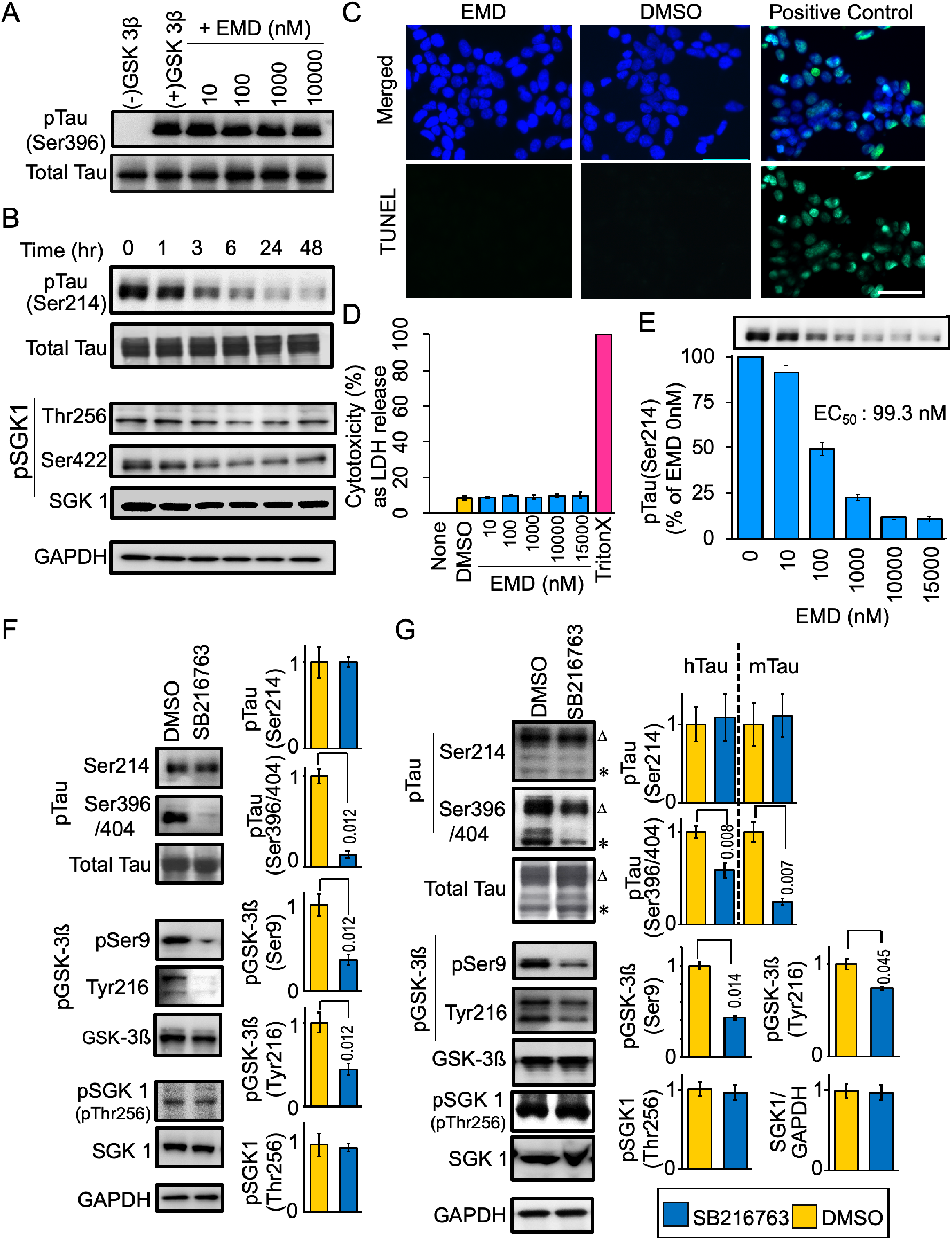
Effect of EMD638683 and SB216763 on tau phosphorylation and cell viability. **(A)** EMD638683 does not directly affect GSK-3ß *in vitro.* Recombinant GSK-3ß was incubated with tau in the presence or absence of EMD638683 and the kinase activity was assessed by tau phosphorylation at Ser396. **(B)** Time-dependent effect of EMD638683 on SGK1 and tau phosphorylation. SH-SY5Y cells stably expressing tau were treated with 1 μM EMD638683 for the indicated periods of time. **(C, D)** EMD638683 does not affect cell viability. **(C)** TUNEL assay of cells treated with 10 μM EMD638683. There were no TUNEL-positive cells (green) after EMD638683 treatment. DNase I-treated SH-SY5Y cells served as a positive control. **(D)** Cytotoxicity was assessed by LDH release in SY-SH5Y cells stably expressing tau 24 hrs after EMD638683 treatment. Triton-X100-treated cells served as a positive control. **(E)** Dose-dependent bar graph for the effect of EMD638683 on pSer214 tau phosphorylation in SH-SY5Y cells. **(F)** GSK-3 specific inhibitor SB216763 inhibits GSK-3ß-associated phosphorylation sites of tau but not SGK1-associated phosphorylation sites. SH-SY5Y cells stably expressing tau were treated with or without 10 μM SB216763 for 48 hrs. Graphs represent quantification of the indicated proteins. DMSO served as a control. **(G)** SB216763 inhibits GSK-3ß-associated phosphorylation sites of tau in the hippocampi of Tg601 mice. SB216763 (5 mg/kg) were stereotactically injected in the Tg601 hippocampi and treated mice were sacrificed after 72 hrs. Graphs represent quantification of the indicated proteins. Scale bar: 100 μm (**C**). Graphs are represented as the means ± SEM (**D, E, F**, n = 5 biological replicates in each group; **G**, n = 3 mice in each group). *p* values are indicated in the graphs. Mann-Whitney U test (**F**) and a paired Student *t*-test (**G**).

**Figure 7.**
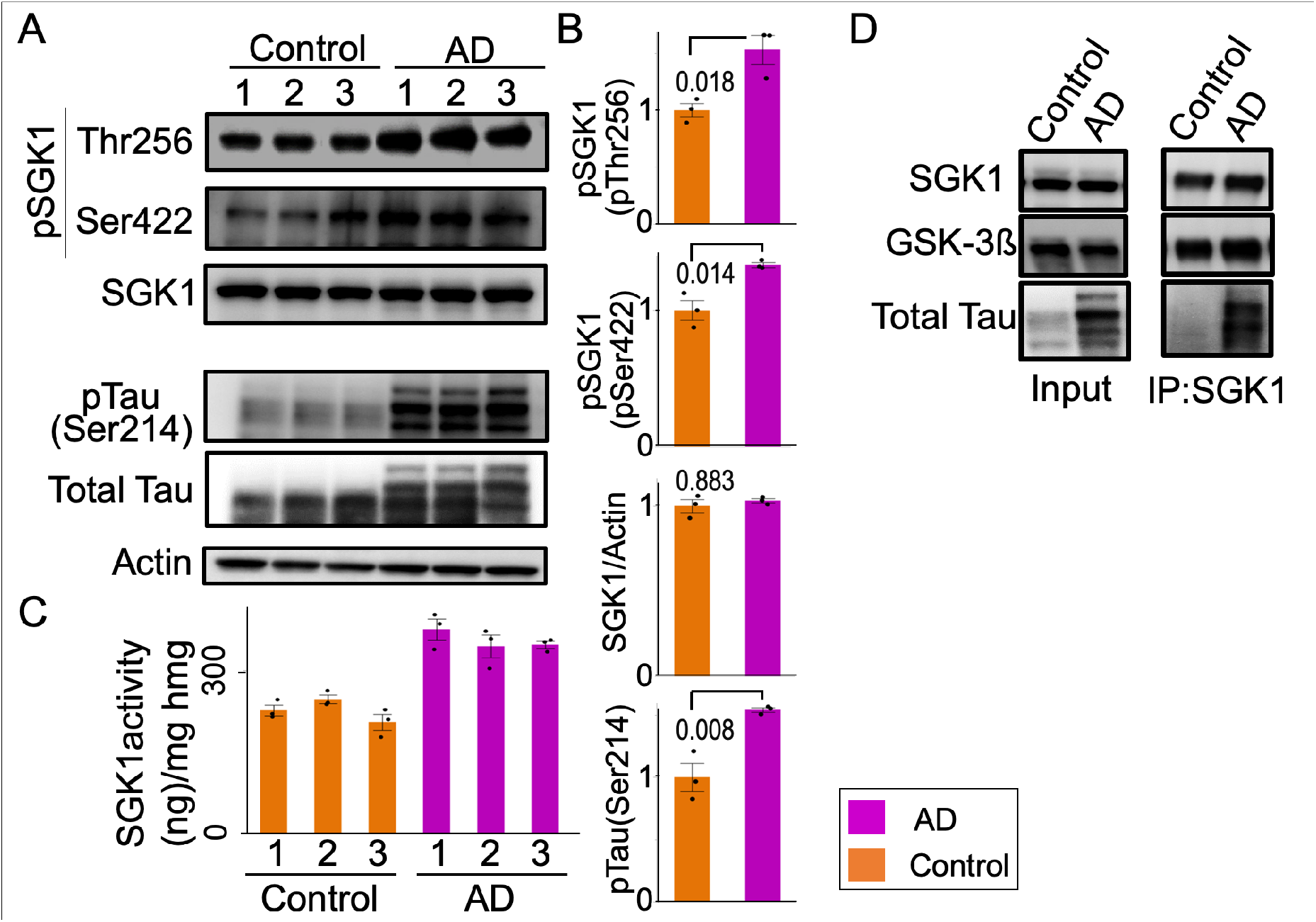
SGK1 is activated in AD brains. **(A, B)** SGK1 phosphorylation and Ser214-phospho-tau levels are increased in the hippocampi of AD brains. Representative Western blots with the indicated antibodies **(A)**. Quantification of the indicated protein levels **(B)**. SGK1, pSGK1 (pThr256 and pSer422), and pSer214-phospho-tau were normalized to actin, SGK1, and HT7 band intensities, respectively. n = 3 individuals. **(C)** Kinase activity of SGK1 immunopurified from human AD brains was measured using an ADP-Glo kinase assay with SGKtide. n = 3 technical replicates. hmg, brain homogenate. **(D)** Co-immunoprecipitation of GSK-3ß and tau with SGK1 from AD brain homogenate using anti-SGK1 antibody. 10% of the homogenate used for immunoprecipitation (IP) for SGK1 and GSK-3ß and 2% for tau were analyzed as input. Representative results were shown from three independent experiments. Graphs in **B, C** represent means ± SEM and *p* values are indicated in the graphs. A paired Student *t*-test **(B)**.

**Figure 7—source data 1.**
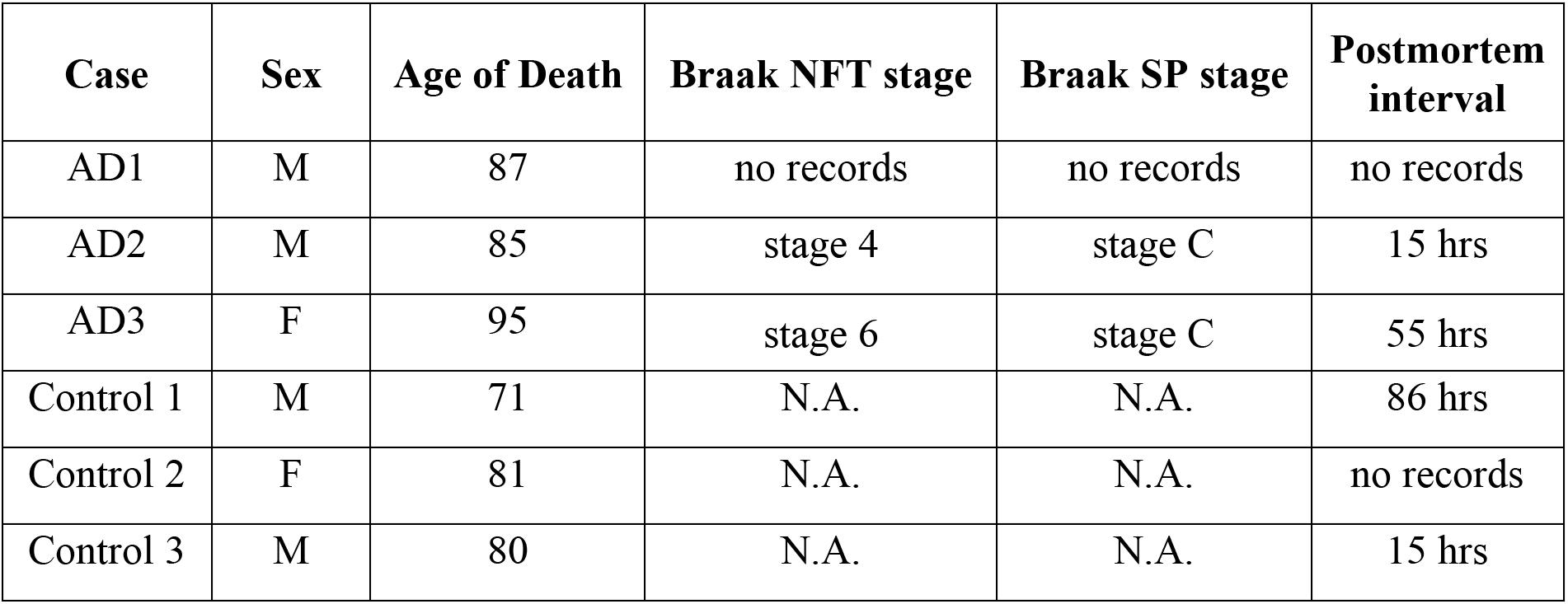
Clinical information on human brain tissues.

## Discussion

Accumulating evidence suggests that obese elderly people exhibit enhanced hippocampal tau phosphorylation (Mrak, 2009). Moreover, T2DM have a 2.5-fold increased risk for cognitive dysfunction (Biessels et al., 2006; Iqbal et al., 2014; Spauwen et al., 2013). The genetic finding of the APOEε4 allele of Apolipoprotein E as the major genetic risk factor for AD also suggest that HFD could modify the AD etiology (Liu et al., 2013). However, the molecular mechanism underlying the acceleration of AD pathology by HFD remains unclear. In this study, we demonstrated that HFD treatment activated SGK1, leading to increased Ser214-phospho-tau levels that resulted in subsequent inhibition of tau’s activity relating to microtubule stabilization in cultured human cells and mice. Moreover, SGK1 activated GSK-3ß through the tertiary SGK1-GSK-3ß-tau complex and induced tau phosphorylation at pSer396 and pSer404, promoting tau fibrilization (Figure 8). Importantly, elevated SGK1 activities and the SGK1-GSK-3ß-tau complex were also found in human AD brain. Moreover, SGK1 inhibitor EMD638683 reduces tau phosphorylation both at Ser214 and Ser396/404, improving the tubulin polymerization activity of tau. Our findings present a concept that SGK1 is a promising molecular target for the prevention of tauopathy in AD.

**Figure 8.**
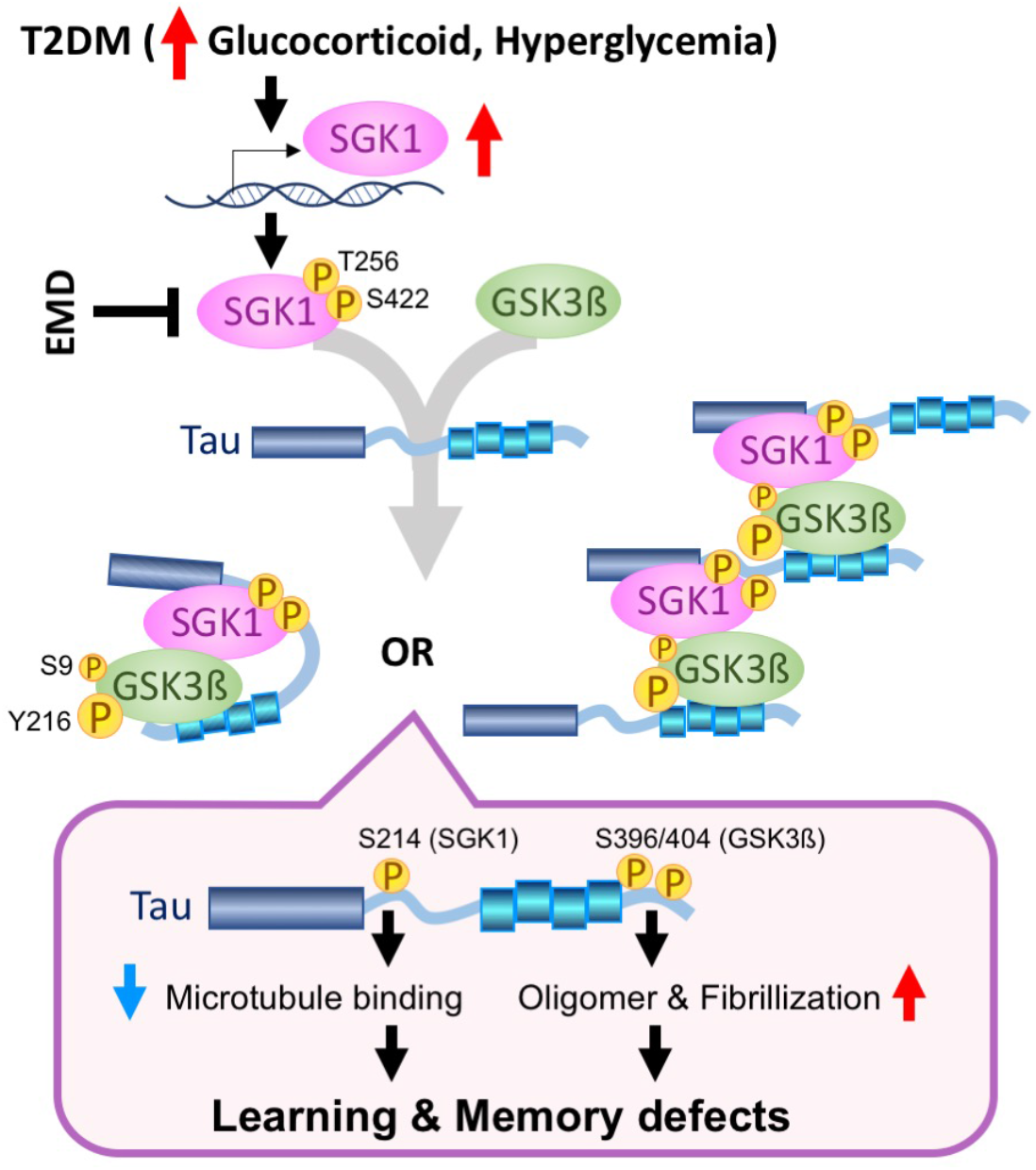
Proposed molecular mechanism for the acceleration of tau pathology by diabetes. Elevated glucocorticoids and hyperglycemia results in transcriptional and post-transcriptional upregulation of SGK1, leading to a tertiary complex containing tau, SGK1, and GSK-3ß. There are two kinds of possible tertiary complexes (left, intramolecular tau complex; right, intermolecular tau complex). The tau tertiary complex promotes GSK-3ß activation and subsequent tau phosphorylation, which results in impaired microtubule binding activity of tau by SGK1-associated phosphorylation at Ser214 and facilitates tau fibrillization by GSK-3ß-associated pSer396/404, leading to defects in learning and memory. A chemical compound, EMD638683 (EMD), inhibits SGK1, thereby suppressing the tertiary complex and tau phosphorylation.

Elevation of serum glucocorticoids is observed in poorly controlled T2DM cases and can be observed in early stages of AD due to malfunction of the hypothalamus-pituitary-adrenal axes (Arnold et al., 2018; Geer et al., 2014; McEwen, 1997). Serum glucocorticoid crosses the blood-brain barrier and stimulates SGK1 expression at the transcriptional level (Mason et al., 2010; Webster et al., 1993). The phenotypes of hyperglycemia and enhanced glucose absorption in *db/db* mice, in which intestinal SGK1 was upregulated, were alleviated by pharmacological inhibition of SGK1 (Li et al., 2016). Pathophysiological involvement of SGK1 in T2DM is strongly suggested by the frequency with which the *SGK1* risk haplotype (I6CC/E8CC/CT) in diabetic patients causes development of obesity, peripheral insulin resistance and inhibition of insulin secretion (Lang et al., 2006; Schwab et al., 2008; Ullrich et al., 2005). Similarly, a molecular relationship between SGK1 activation and tau modification has been suggested (Green et al., 2006; Lang et al., 2006). Overexpression of TGF-ß, which is a potent stimulator of SGK1, increased phosphorylated tau in mouse brain tissues (Lang et al., 2006). Intraperitoneal injection of DXM elevated tau phosphorylation and altered the hippocampal cognitive function in 3X-Tg-AD mice (Green et al., 2006). Consistent with these studies, SGK1 activation and subsequent tau phosphorylation at pSer214 in the *db/db* T2DM genetic model mice were observed, while exogenous leptin might also contribute to cognitive impairment (Dey et al., 2017). Enhanced SGK1 activation and elevated Ser214-phospho-tau in the hippocampus were also observed in an STZ-treated mouse model of diabetes (Figure 2-figure supplement 4). Reduction in body temperature has been suggested as a mechanism for tau phosphorylation in STZ-treated or leptin receptor-deficient mice (Gratuze et al., 2017; Planel et al., 2007). While we observed that *db/db*, HFD-treated, or STZ-treated mice have common phenotypes of increased SGK1 activation, elevated glucocorticoids and hyperglycemia, impaired thermoregulation was not observed in HFD-treated mice (Figure 1-figure supplement 1G). Therefore, hypothermia would not be involved in the elevated tau phosphorylation observed in these diabetic models.

A previous study reported that transgenic mice expressing human AD-associated tau G272V/P301S double mutations exhibited insulin resistance similar to non-transgenic littermates, while the spatial learning impairments by mutant tau was promoted by HFD (Leboucher et al., 2013). In contrast, the current study indicated that the behavioral disturbances by HFD treatment were similar in both Tg601 and NTg mice. Our Tg601 mice, which express wild-type human tau under the CaMKIIα promoter, exhibit no apparent AD-like behavioral phenotypes, but an obvious tauopathy and mild defects in the hippocampal activities (Hara et al., 2017; Kambe et al., 2011). Our results suggest that HFD-induced tau hyperphosphorylation is a stronger risk than tau protein level at least under the condition of wild-type tau.

Elevated glucocorticoid levels and mild hyperglycemia were observed in HFD-treated mice, as previously reported in type 2 and type 1 diabetes models (Revsin et al., 2008; Stranahan et al., 2008). Moreover, HFD-treated mice develop lipid mediated insulin resistance in the brain (Kothari et al., 2017). Considering three conditions associated with T2DM, we evaluated SGK1 activity in cultured cells treated with DXM, high glucose or PA-BSA. Among these 3 conditions, DXM and high glucose exposure upregulated and activated SGK1, leading to increased tau phosphorylation, in cultured cells, and the effects of DXM were stronger than those of high glucose treatment (Figure 4). SGK1 is involved in the upregulation of sodiumglucose cotransporter 1 (SGLT1) and enhances intestinal glucose absorption, ultimately leading to obesity, elevated circulating insulin and insulin resistance in the peripheral tissues (Lang et al., 2006). Moreover, the elevated peripheral insulin concentration might be associated with the reduction in brain insulin production or uptake reported in AD brains and might lead to insulin resistance (Blázquez et al., 2014). Although the effect of insulin resistance in the central nervous system on tau phosphorylation is a matter of debate, insulin resistance has been demonstrated to exacerbate Aß deposition by decreasing levels of insulin degradation enzyme, which is a key enzyme for Aß degradation, and/or by increasing γ-secretase activity that causes increased Aß production (Mullins et al., 2017). While PA-BSA treatment in cultured cells did not upregulate and activate SGK1, increased Ser396/Ser404-phospho tau was still observed (Figure 4A). Considering the downregulation of pSer473 AKT and pSer9 GSK-3ß by PA-BSA, hyperlipidemia could activate GSK-3ß via inactivation of the AKT pathway. However, these molecular changes by PA-BSA were not observed in the hippocampi of HFD-treated mice (Figure 1D *vs.* Figure 4-figure supplement 1A).

The activity of GSK-3ß is positively and negatively regulated by Tyr216 and Ser9 phosphorylation, respectively. We observed the elevated phosphorylation at both sites in HFD-treated mice, *db/db* mice and STZ-treated mice whereas pSer9 phosphorylation was not altered in SGK1-knockdown SH-SY5Y cells, suggesting that pSer9, but not pTyr216, is regulated by different kinases other than SGK1. A recent report demonstrated that pSer9 is a minor phosphorylation site of GSK-3ß when compared with pTyr216 (Krishnankutty et al., 2017), suggesting that the contribution of pTyr216 is more important to GSK-3ß kinase activity.

We demonstrated that tau phosphorylation at Ser214 by SGK1 drastically reduced the tubulin polymerization abilities of tau. Thus, SGK1 might directly affect microtubule stabilization by disassembling microtubules (Yang et al., 2006). Hyperphosphorylation of tau in the proline-rich domain or the microtubule-binding repeat is associated with inhibition of microtubule assembly, whereas C-terminal phosphorylation in tau was mostly involved in tau self-aggregation (Liu et al., 2007). Additionally, the literature describes that tau phosphorylation at Ser214 primes GSK-3ß-mediated tau phosphorylation at the C-terminal region, which is consistent with our findings (Figure 5-figure supplement 2B) (Liu et al., 2004). In addition, pSer214 phosphorylation by SGK1 promoted the interaction of tau with 14-3-3 proteins, which might increase the aggregation propensity of tau *in vivo* (Chun et al., 2004; Sadik et al., 2009). Thus, our data showing tau phosphorylation at Ser214 by SGK1 and the accompanying tau pathology and impaired cognitive functions in mice corroborate the previous findings (Liu et al., 2007). Furthermore, our study revealed a complex molecular mechanism underlying SGK1-mediated activation of GSK-3ß and tau phosphorylation at the C-terminal region, promoting tau aggregation. In the absence of tau, SGK1 negatively regulated GSK-3ß kinase activity by increasing phosphorylation at Ser9, which was observed with CTF tau and GSK-3ß substrate peptide. However, in the presence of full-length tau or NTF, phosphorylation of GSK-3ß at Tyr216 was increased. The increase in Tyr216 phosphorylation might be an effect of autophosphorylation or an intramolecular process related to a conformational change of GSK-3ß (Cole et al., 2004). The binding of the SGK1-GSK-3ß complex to the N-terminal region of tau facilitates GSK-3ß phosphorylation at Tyr216 and resultant activation of GSK-3ß, ultimately leading to tau hyperphosphorylation. This substrate-dependent activation of GSK-3ß was also observed in Ser214Ala-mutated tau, suggesting that a priming effect of Ser214 phosphorylation by SGK1 has a limited impact (Figure 5D and E). Another site of tau phosphorylated by SGK1, which was detected by phos-tag Western blot, might also contribute to GSK-3ß activation (Figure 5-figure supplement 2C).

This study also raises the possibility of the limitation of GSK-3ß-targeted therapeutics in terms of tauopathy, while SGK1 could be a promising target for affecting the pathogenesis of AD. SGK1 appeared to phosphorylate tau at Ser214 before affecting the phosphorylation of GSK-3ß at Tyr216, suggesting that the dissociation of tau from the microtubule is an earlier event than GSK-3ß activation and subsequent tau phosphorylation by GSK-3ß (Figure 4-figure supplement 1C). Thus, SGK1 would be a more effective target than GSK-3ß. Moreover, our hypothesis was supported by the ineffectiveness of GSK-3ß inhibitor SB-216763 in reducing pSer214-Tau (Figure 6-figure supplement 1F and G).

Abundant expression of SGK1 has been previously reported in the AD brain (Sahin et al., 2013). Although SGK1 protein levels in the hippocampi of AD cases were not different from those in healthy controls, the active form of SGK1 and Ser214-phospho-tau levels were increased (Figure 7A-C). Moreover, the existence of the SGK1-GSK-3ß-tau tertiary complex in the AD brain reinforced the idea of T2DM and AD overlapping through corticosteroid effects and indicated the molecular link for a common therapeutic approach. Pharmacological inhibition of SGK1 by EMD638683 effectively reduced the fasting blood glucose and its biomarker HbA1c because SGK1 positively regulates SGLT1 (Li et al., 2016). Our study first showed that inhibiting SGK1 by EMD638683 suppresses GSK-3ß activity and tau phosphorylation, although no direct inhibitory activity of EMD638683 on GSK-3ß was observed (Figure 8 and Figure 6-figure supplement 1A).

Previous studies have demonstrated that tau hyperphosphorylation in HFD-treated mice is caused by dysregulation of insulin signaling, supporting the finding that patients demonstrating an elevated fasting glucose are at an increased risk for AD (Morris et al., 2016). We revealed that increased glucocorticoid levels and hyperglycemia observed in T2DM activate SGK1, which phosphorylates tau directly at Ser214 and indirectly at Ser396/Ser404 by promoting increased GSK-3ß activation. Given that SGK1 is involved in the two important events in AD etiology, tau phosphorylation and GSK-3ß activation, SGK1 would be a promising target for AD and T2DM therapies.

## Materials and Methods

### Key resources table

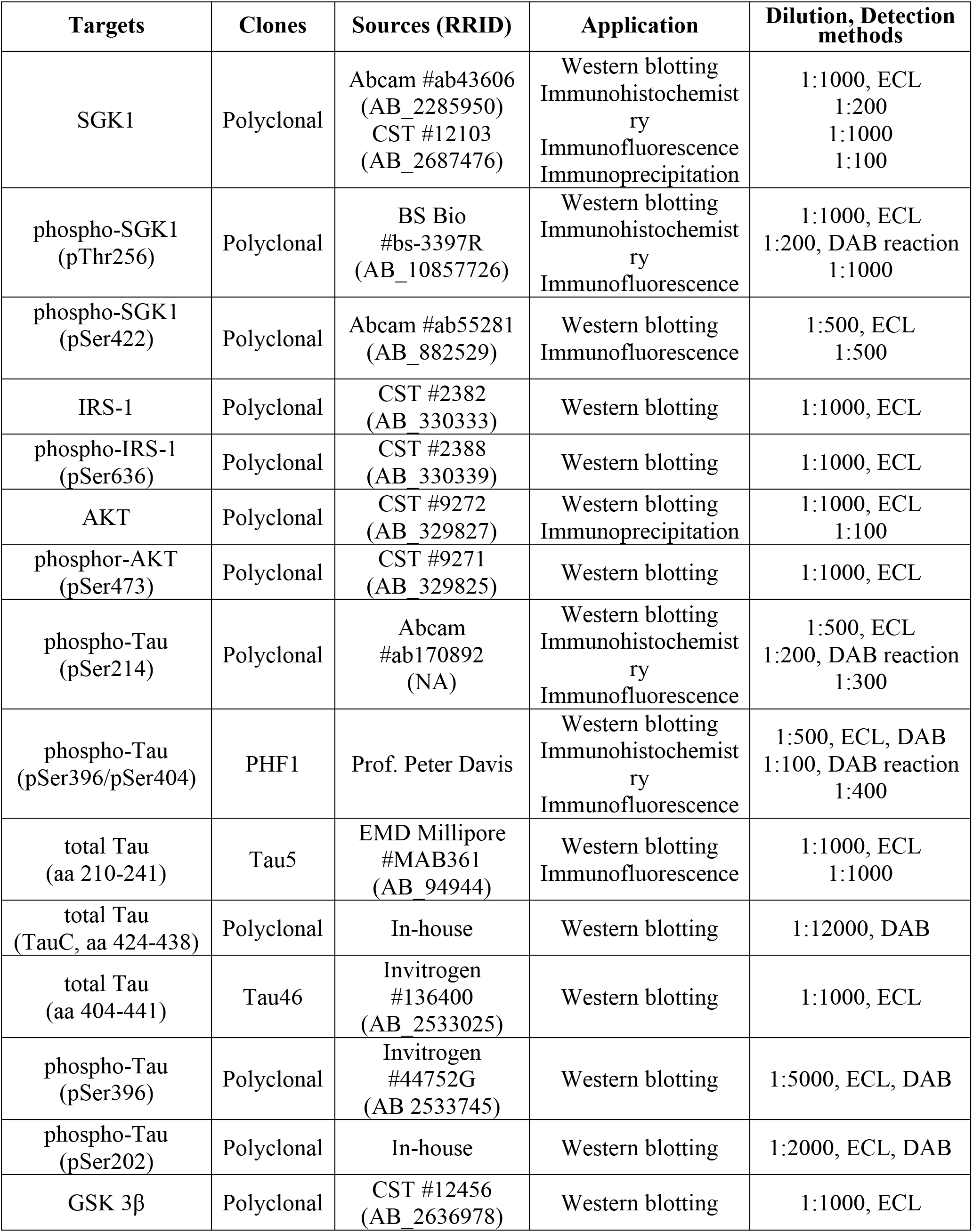
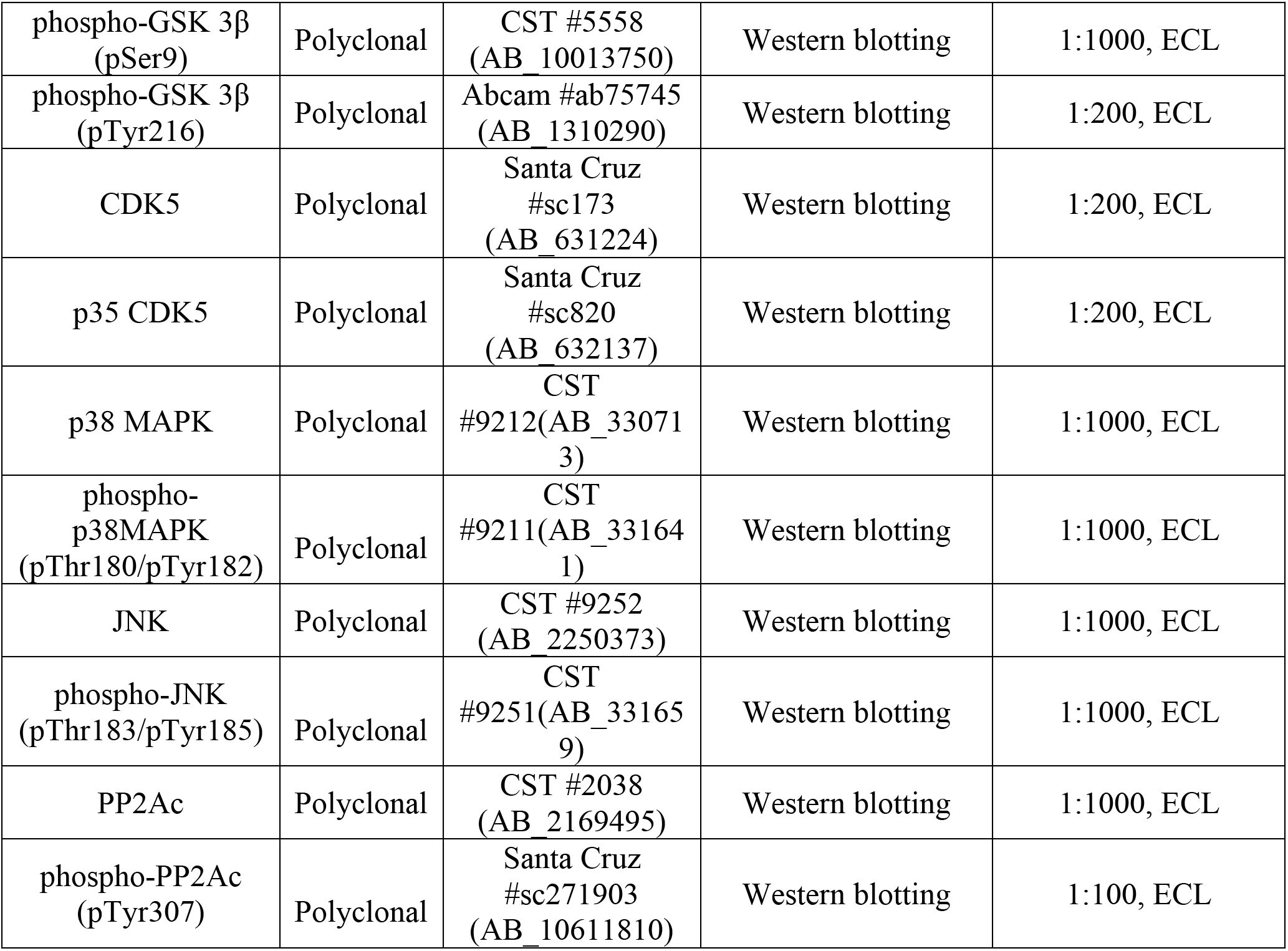

| Targets | Clones | Sources (RRID) | Application | Dilution, Detection methods |
| --- | --- | --- | --- | --- |
| SGK1 | Polyclonal | Abcam #ab43606 (AB_2285950)<br>CST #12103 (AB_2687476) | Western blotting<br>Immunohistochemistry<br>Immunofluorescence<br>Immunoprecipitation | 1:1000, ECL<br>1:200<br>1:1000<br>1:100 |
| phospho-SGK1 (pThr256) | Polyclonal | BS Bio #bs-3397R (AB_10857726) | Western blotting<br>Immunohistochemistry<br>Immunofluorescence | 1:1000, ECL<br>1:200, DAB reaction<br>1:1000 |
| phospho-SGK1 (pSer422) | Polyclonal | Abcam #ab55281 (AB_882529) | Western blotting<br>Immunofluorescence | 1:500, ECL<br>1:500 |
| IRS-1 | Polyclonal | CST #2382 (AB_330333) | Western blotting | 1:1000, ECL |
| phospho-IRS-1 (pSer636) | Polyclonal | CST #2388 (AB_330339) | Western blotting | 1:1000, ECL |
| AKT | Polyclonal | CST #9272 (AB_329827) | Western blotting<br>Immunoprecipitation | 1:1000, ECL<br>1:100 |
| phosphor-AKT (pSer473) | Polyclonal | CST #9271 (AB_329825) | Western blotting | 1:1000, ECL |
| phospho-Tau (pSer214) | Polyclonal | Abcam #ab170892 (NA) | Western blotting<br>Immunohistochemistry<br>Immunofluorescence | 1:500, ECL<br>1:200, DAB reaction<br>1:300 |
| phospho-Tau (pSer396/pSer404) | PHF1 | Prof. Peter Davis | Western blotting<br>Immunohistochemistry<br>Immunofluorescence | 1:500, ECL, DAB<br>1:100, DAB reaction<br>1:400 |
| total Tau (aa 210-241) | Tau5 | EMD Millipore #MAB361 (AB_94944) | Western blotting<br>Immunofluorescence | 1:1000, ECL<br>1:1000 |
| total Tau (TauC, aa 424-438) | Polyclonal | In-house | Western blotting | 1:12000, DAB |
| total Tau (aa 404-441) | Tau46 | Invitrogen #136400 (AB_2533025) | Western blotting | 1:1000, ECL |
| phospho-Tau (pSer396) | Polyclonal | Invitrogen #44752G (AB_2533745) | Western blotting | 1:5000, ECL, DAB |
| phospho-Tau (pSer202) | Polyclonal | In-house | Western blotting | 1:2000, ECL, DAB |
| GSK 3β | Polyclonal | CST #12456 (AB_2636978) | Western blotting | 1:1000, ECL |
| phospho-GSK 3β<br>(pSer9) | Polyclonal | CST #5558<br>(AB_10013750) | Western blotting | 1:1000, ECL |
| phospho-GSK 3β<br>(pTyr216) | Polyclonal | Abcam #ab75745<br>(AB_1310290) | Western blotting | 1:200, ECL |
| CDK5 | Polyclonal | Santa Cruz<br>#sc173<br>(AB_631224) | Western blotting | 1:200, ECL |
| p35 CDK5 | Polyclonal | Santa Cruz<br>#sc820<br>(AB_632137) | Western blotting | 1:200, ECL |
| p38 MAPK | Polyclonal | CST<br>#9212(AB_33071<br>3) | Western blotting | 1:1000, ECL |
| phospho-<br>p38MAPK<br>(pThr180/pTyr182) | Polyclonal | CST<br>#9211(AB_33164<br>1) | Western blotting | 1:1000, ECL |
| JNK | Polyclonal | CST #9252<br>(AB_2250373) | Western blotting | 1:1000, ECL |
| phospho-JNK<br>(pThr183/pTyr185) | Polyclonal | CST<br>#9251(AB_33165<br>9) | Western blotting | 1:1000, ECL |
| PP2Ac | Polyclonal | CST #2038<br>(AB_2169495) | Western blotting | 1:1000, ECL |
| phospho-PP2Ac<br>(pTyr307) | Polyclonal | Santa Cruz<br>#sc271903<br>(AB_10611810) | Western blotting | 1:100, ECL |

### Animals and diet

Tau transgenic model (Tg601) mice overexpressing wild-type human tau (2N4R) under the control of the CAMKIIα promoter were developed in our laboratory (Kambe et al., 2011). Tg601 mice and their littermate non-transgenic (NTg) mice have the same C57BL/6J genetic background. All animal experiments were performed in accordance with the regulations outlined by Japanese law and the National Institutes of Health guidelines and were also approved by the Animal Care and Experimentation Committee of Juntendo University. Ten-month-old female Tg601 and NTg mice were separated into HFD and CD groups (n = 27 in each group, respectively). The HFD group was fed a high-fat diet (D12492, Research diet Inc, USA) containing 35 wt% fat, 26 wt% carbohydrate, and 26 wt% protein with an additional 5 wt% sucrose and the CD groups were fed with control chow (CRF–1, Oriental Yeast Co. Ltd., Japan) containing 5.4 wt% fat, 53.8 wt% carbohydrate, 21.9 wt% protein for 5 months. Fiber and mineral contents were not exactly controlled between HFD and CD.

### *db/db* mice

Female B6 Leprdb/J (*db/db*) homozygous mice were obtained from Jackson Laboratories at 5 weeks of age. On arrival to the animal facility, mice were housed 3 per cage under the SPF condition. Mice were sacrificed at 4 months of age.

### STZ injection and preparation of *in vivo* hyperglycemia

*In vivo* hyperglycemia was developed as our previous report (Elahi et al., 2016b). Briefly, STZ (60 mg/kg body weight) was injected into Tg601 female mice (n = 6, 12.0 ± 2.0 months old) and NTg female mice (n = 6, 12.0 ± 1.0 months old) for 5 consecutive days. Control female mice (Tg601 mice, n = 6; NTg mice, n = 6) were injected with 0.9% saline (Otsuka, Japan). Blood glucose levels were assessed at 0, 4, 7, 14, and 21 days after the STZ injection.

### Mice sacrifice

Mice used for biochemical and RNA analysis were sacrificed by cervical dislocation after behavioral assessments to obtain the hippocampus for RNA extraction, SGK1 kinase assay and glucocorticoid measurement. The left and right sides of hippocampi were subjected to histochemical analysis and biochemical analyses, respectively.

### Glucose and insulin tolerance tests

The intraperitoneal glucose tolerance test (IPGTT) and intraperitoneal insulin tolerance test (ITT) were carried out to confirm whether HFD-treated mice developed T2DM-like features. After 2 months of HFD treatment, mice were subjected to IPGTT and ITT. IPGTT was carried out by injecting D (+) glucose (2 g/kg body weight) intraperitoneally following 8 hrs fasting. ITT was performed following a 6-hr fast with intraperitoneal insulin injection (0.75 U/kg). Blood glucose was measured at 0, 15, 30, 60, 90, and 120 min following injections by collecting blood from the tail vein using a One Touch Lifescan-LFS ultra-glucometer (LifeScan Inc., USA).

### Behavioral analyses

Mice were subjected to 4 behavioral batteries 5 months after HFD treatment. To assess spatial learning, Morris water maze tests, mnemonic function, and the Y-maze test were performed. The elevated plus maze test and rotarod test were assessed to evaluate anxiety and motor function, respectively.

### Morris water maze

To measure reference learning (acquisition) and memory (retention), mice were placed into a 122-cm water-filled circular pool. The pool was divided into four equal-sized quadrants and was surrounded by various visual cues, including a beach ball and a colored wall poster with a specific sign, which helped the mice to orient their location in the pool. A clear, 9-cm platform was placed in the middle of quadrant II. The surface of the platform was exactly 1 cm below the water surface and visually indiscernible to the mice. Reference learning (acquisition) was evaluated over a 5-day period, with three successive sessions per day. The swimming path, swimming time, and latency to find the platform (up to 60 s) for each trial were recorded using an automated video tracking system (Muromachi Kikai, Japan).

### Y-maze

As a measure of general activity and short-term working memory function, mice were tested in a Y-maze with three arms (identified as A, B, and C). Each mouse was placed into one of the three arms facing the middle area, and allowed to explore the maze for 8 min. The total number of arm entries and sequence of arm choices were observed and recorded. Alternation was defined as the ratio of arm choices differing from the previous two choices divided by the total number of entries. The alternation score was expressed as a percentage.

### Elevated plus maze

Anxiety/emotionality were assessed by using a plus-shaped maze elevated 82 cm above the floor (Muromachi Kikai). The maze consists of four arms, each 30 cm, including two opposite “closed” arms surrounded by dark walls and two opposite “open” arms that are exposed without any walls. Each mouse was placed at the center of the maze facing a closed arm, and allowed to freely explore the maze for a period of 8 min. During this trial, the amount of time (in seconds) spent in the open arms was observed and recorded using an automated video tracking system (Muromachi Kikai).

### Novel object recognition test

Novel object recognition test was used for assessment of learning and memory in rodents (Webster et al., 2013). A cylindrical plastic box of 50 × 50 × 50 cm (Muromachi Kikai) was installed to create an open field in the test. Objects that differ in shape and texture were used as test objects. On a training day, mice were familiarized with two identical objects. On a testing day, a different object (novel object) was replaced with one familiar object. The exploratory time with a novel object during the test session was expressed as recognition index (RI). RI (%) was calculated as exploratory time with a novel object / (exploratory time with a novel object + exploratory time with a familiar object) × 100.

### Rotarod

Motor neuronal activity was assessed using the rotarod performance test. Locomotor behavior analysis was carried out 5 months after HFD treatment using a rotarod machine with automatic timers and falling sensors (MK-660D, Muromachi Kikai). Mice were placed on a 3-cm diameter rotating rod covered with rubber, and rotation was accelerated from 3 to 35 rpm over 5 min. Fall latency was recorded for each trial.

### Antibodies

All primary antibodies used are listed in **Key resources table**.

### RNA isolation, microarray and reverse transcription-PCR

Hippocampal total RNA was isolated using an Agilent total RNA isolation mini kit following the manufacturers’ instructions (Agilent, USA). The amounts of RNA were quantified by a nanodrop spectrophotometer (Thermo Fisher Scientific, USA), and the RNA integrity and quality were assessed by an Experion automated electrophoresis system (Biorad, USA). Expression profiles of the RNA samples with an RQI ≥ 8.0 and a 28S/18S rRNA ratio ≥ 1 in Experion analysis were determined by using Affymetrix GeneChip Mouse transcriptome assay 1.0 (MTA 1.0) arrays (Thermo Fisher Scientific) according to the manufacturer’s instructions. The GeneChip WT Expression Kit and GeneChip WT Terminal Labeling and Controls Kit (Thermo Fisher Scientific) were used to generate amplified and biotinylated sense-strand cDNA from expressed transcripts (250 ng of total RNA). Manufacturer’s instructions were followed for the hybridization, washing, and scanning steps. The CEL files generated during scanning were processed by Expression console software (Thermo Fisher Scientific) to generate CHP files. The CHP files confirmed the data quality and were further analyzed by Transcriptome array console (TAC, ver. 3.1, Applied Biosystems) to obtain gene-level and exon-level estimates and alternative splicing information for all transcripts. The gene-level estimates were subjected to statistical analysis and hierarchical and partitioning clustering by TAC, which also link with integrated pathway analysis.

Reverse transcription of the *sgk1* gene was carried out by synthesizing complementary DNA (cDNA) from total RNA transcripts. M-MLV Reverse Transcriptase (Promega) and a set of random hexamers were used with 1 μg total RNA following the manufacturer’s instructions. Amplification of the *sgk1* and *gapdh* genes was carried out by KOD-PLUS-ver. 2 (Toyobo Life Science, Japan) using primers designed with the following sequences: *sgk1*, forward 5’-CGTCAAAGCCGAGGCTGCTCGAAGC-3’ and reverse 5’-GGTTTGGCGTGAGGGTTGGAGGAC-3 and for *gapdh*, forward 5’-TCACCACCATGGAGAAGGC-3’ and reverse 5’-GCTAAGCAGTTGGTGGTGCA-3’, respectively.

### Preparation of AAV-mSGK1 and stereotactic surgery

Recombinant AAV, pAAV2-SynTetOff-mSGK1-2A-EGFP, was prepared by replacement of GFP sequence in pAAV2-SynTetOff-GFP (Sohn et al., 2017) with mSGK1-2A-EGFP. pAAV2-SynTetOff-mSGK1-2A-EGFP, pHelper, and pBSIISK-R2C1 were co-transfected into HEK293T cells for the virus production. Virus particles were purified from the cells after 72 hrs of medium replacement. Following the extraction by three freeze-and-thaw cycles, the virus particles were recovered from post-ultracentrifugation supernatant and concentrated by ultrafiltration. The virus titer was determined by qPCR for AAV2 inverted terminal repeat sequences. AAV2/1-SynTetOff-mSGK1-2A-EGFP (1.08 × 10^14^ vg in 2.5 μL) were injected into the mouse hippocampus through stereotactic surgery.

### Mouse brain tissue and cell preparation for biochemical analysis

Frozen hippocampi were homogenized and sonicated in 10X volume of homogenization buffer (HB) containing 10 mM Tris–HCl, pH 7.5, 0.8 M NaCl, 1 mM EGTA, 1 mM dithiothreitol (DTT), and 0.1 μM phenylmethanesulfonylfluoride. The supernatants obtained following 20,000 g centrifugation for 20 min at 4°C were used for further analysis. Samples for Western blotting were prepared after protein standardization using a pierce BCA assay (Thermo Fisher Scientific) using a 2X volume of loading buffer.

To prepare heat-stable tau, 1 mM DTT and 1% ß-mercaptoethanol were added to the supernatant and were exposed to heat denaturation for 5 min at 100°C. The samples were centrifuged at 25,000 g for 20 mins, and then, the supernatant was collected. The supernatants were dialyzed against 30 mM Tris-HCl, pH 8.5, and concentrated using an Amicon filter device. Detergent-soluble and -insoluble tau fractions were prepared from the frozen hippocampus from the right half of the brain following the previously described protocol (Elahi et al., 2016b).

Briefly, after sonication, 100 μl homogenates were centrifuged at 100,000 x g for 20 min at 4°C. The supernatants (TS: Tris soluble) were collected, while the pellets were resuspended in 100 μl of HB containing 1% (v/v) Triton X100 and incubated for 30 min at 37°C. After centrifugation for 20 min at 100,000 x g, the Triton X100-soluble fractions (TX) were collected, and the pellets were further resuspended in 50 μl of HB containing 1% sarkosyl. The samples were centrifuged, and the supernatants were collected as the sarkosyl-soluble fractions (SS). The resulting pellets were used as the sarkosyl-insoluble fraction (Ppt) for further analysis. The same fractionation protocol was applied to SH-SY5Y cells expressing tau.

### Western blotting analysis

Western blotting samples were separated by SDS-PAGE SuperSep 5-20% (Wako, Japan) and were blotted onto polyvinylidene membranes (Immobilon-P). The blots were incubated at 4°C or room temperature overnight with a primary antibody, followed by interactions with an appropriate and conjugated secondary antibody after 1 hr blocking with skim milk or 1% gelatin. The targeted protein bands were visualized with extended chemiluminescent (ECL) reaction using Immobilon western HRP substrate (Millipore) or by the VECTASTAIN ABC Kit (VECTOR Laboratories, USA) and 3,3’-diaminobenzidine (DAB) containing NiCl2 substrate. Band intensities were analyzed using ImageJ software (National Institutes of Health, USA).

### Brain autopsy

Brain tissues from three normal elderly subjects as healthy controls and three pathologically confirmed AD patients (Braak NFT stage 4-6) were obtained from Juntendo University hospital brain bank. All experimental procedures for brain autopsy were approved by the Juntendo University School of Medicine Ethics Committee (Approval number:2012068). Details of the brain samples are described in Figure 7—source data 1. Hippocampi were homogenized with a polytron homogenizer in HB buffer containing 10% sucrose. The homogenate was centrifuged for 20 min at 20,000 g and the supernatant was retained. The pellet was further solubilized with half volume buffer and recentrifuged. Both supernatants were combined for further analysis.

### Glucocorticoid ELISA

Mouse brain glucocorticoid levels were measured using a mouse endogenous glucocorticoid (E-GC) ELISA kit (Blue gene biotechnology, Shanghai, China) after extraction of total corticosteroid from brain homogenate. Brain homogenate was prepared in 1X PBS containing complete protease inhibitor cocktail (Roche). Total corticosteroid was extracted by mixing equal volume of brain homogenate with methanol: chloroform (2:1) solution. After careful mixing and centrifugation, the chloroform layer was removed to a new tube, where then the solvent was evaporated. The ppt was dissolved in 0.1 ml PBS and applied to a precoated 96-well ELISA plate. We followed further instructions provided by the supplier for measurement. The plates were analyzed using SpectraMax iD3 Multi-Mode Microplate Readers (Molecular Device). The data were expressed as ng of glucocorticoid/mg of brain homogenate.

### Immunohistochemistry and immunocytochemistry

Immunohistochemistry was carried out by preparing paraffin-embedded sections and following the previously described method (Elahi et al., 2016a). Briefly, five-micrometer-thick coronal sections were deparaffinized and rehydrated in a series of xylene and graded ethanol solutions. Endogenous peroxidase was quenched by treating with 0.3% H_2_O_2_. Antigen retrieval was carried out by autoclaving at 120°C for 5 min. The primary antibody was applied overnight at 4°C, followed by specific secondary immunoglobulin coupled with HRP (Histofine simple stain MaxPo M, Nichirei Bioscience Inc., Japan) for a 60-min incubation at room temperature. All antibody dilutions and washing steps were performed in phosphate buffer, pH 7.2. HRP intensities in the hippocampus including the CA1, CA3, CA4, and dentate gyrus regions in three sections/slices from each group of mice were measured using the ImageJ Fiji platform. Data were represented as staining intensity per square millimeter.

For immunofluorescence staining, cells were cultured on a collagen-coated coverslip. After specific treatments, cells were fixed with 4% paraformaldehyde-PBS. The cell membrane was permeabilized with 5 μg/ml digitonin followed by blocking with 5% BSA. After probing with primary antibodies, specific Alexa-fluor labeled secondary antibody were used. 4’,6-Diamidino-2-phenylindole dihydrochloride (DAPI) staining was used to visualize the nuclei (Thermo Fisher Scientific). Fluorescence was assessed using a fluorescence laser microscope (LSM780, Zeiss, Germany).

### Establishment of SH-SY5Y cells stably expressing 2N4R tau

SH-SY5Y cells were cultured in DMEM/F-12 medium (Sigma-Aldrich) with 10% fetal bovine serum at 37°C under a humidified atmosphere of 5% CO_2_. Cells were plated on six-well culture plates at 50% confluency and then transfected with 1 μg/well of the expression plasmid (pCDNA3.1-hTau441) using X-tremeGENE9 DNA transfection reagent (Roche). Cells were transferred to a 10-cm dish containing 300 μg/ml gentamycin-disulfate (G418, Nacalai tesque) selection media 24 hrs post-transfection. After one month of selection, the cells were plated onto a 96-well plate using a cell sorter. Pure single clones were selected by confirming the expression of tau, NTF or CTF expression via Western blotting.

### Palmitic acid-BSA conjugate preparation

Palmitic acid (PA) was dissolved and saponified in 0.1 M KOH in ethanol and prepared into a 500 mM solution at 70°C. We prepared 10% fatty-acid-free BSA in DMEM, and a final concentration of 5 mM PA was added to 1 ml BSA, dissolving by vortexing and allowing a heating-sonication-heating cycle (15 min). The resultant solution was incubated at 55°C for 15 min, sonicated for 15 min in a water sonicator and further incubated at 55°C for 15 min. A vehicle control was prepared by dissolving 10 μl 0.1 M KOH/Ethanol in BSA. PA-BSA was stored at −30°C and defrosted and melted in a water bath at 55°C for 15 min before use. The molar ratio of the PA-BSA conjugate was 3:1 (PA: BSA) and 5 μM and 50 μM of the PA-BSA conjugates were used for treatment.

### DXM, high glucose and palmitic acid treatments

For treatment with DXM or glucose, cells were transferred to 6-well plates and allowed to grow at over 70% confluency for 48 hrs. Forty-eight hours before the experiments, the cells were transferred to DMEM containing 5.5 mM glucose, which is a physiological concentration of blood glucose, and 2% serum replacement (Thermo Fisher Scientific). DXM stock solution was prepared in a DMSO: ethanol (1:1) solution. Glucose was added to the cells to reach a final concentration of 10 to 50 mM, mimicking mild to typical hyperglycemia (final concentrations in media: 10 mM, 180 mg/dl; 20 mM, 360 mg/dl; 33.3 mM, 600 mg/dl; 50 mM, 900 mg/dl). Similar mannitol concentrations were used to control for changes in osmolarity. To evaluate the effect of fatty acid-mediated insulin resistance, cells were treated with palmitic acid-BSA conjugate for 48 hrs as described previously (Kim et al., 2017).

### siRNA transfection

siRNA transfection was carried out using X-tremeGENE siRNA transfection reagent (Roche) at 70% cell confluency. Medium was replaced with fresh medium 24 hrs post-transfection and maintained for another 24 hrs. The sequence of siRNA against SGK1 was sense, 5’-GUCCUUCUCAGCAAAUCAA-3’ and antisense, 5’-UUGAUUUGCUGAGAAGGAC-3’ (Thermo Fisher Scientific).

### Recombinant tau preparation and *in vitro* kinase assay

A bacterial expression plasmid pRK172 harboring human tau, N-terminal (1-242) tau/NTF or N-terminal (243-441) tau/CTF cDNA was used to produce recombinant tau protein in *Escherichia coli* BL21 (DE3) (Merck Millipore). Purification was performed as described previously (Hasegawa et al., 1997). Briefly, harvested bacterial pellets were lysed by sonication using tau purification buffer containing 50 mM PIPES, 5 mM EGTA, and 1 mM DTT. The supernatant was subjected to heat treatment for 5 min in the presence of 1% ß-mercaptoethanol. The resultant clear supernatant was passed through a Q Sepharose column, and tau protein was eluted against 350 mM NaCl and dialyzed against 50 mM Tris–HCl, pH 7.5. Recombinant SGK1 and GSK-3ß were purchased from SignalChem (Richmond, Canada).

The tau phosphorylation assay was carried out at 30°C in 30 μl of kinase buffer containing 30 mM HEPES, 10 mM MgCl2, 400 μM ATP, 10 mM DTT, 10 mM ß-glycerophosphate and 5 μg tau. Recombinant SGK1 or GSK-3ß was added to initiate the reaction, and the reaction was terminated by heat inactivation. In some experiments, two protein kinases were used in combination or added sequentially, and tau phosphorylation was detected by Western blotting with specific antibodies. Other substrates (2N4R tau with Ser214Ala mutation, NTF and CTF tau or SGKtide or GSK-3ßtides) in the kinase reaction were also applied in the same amount and the kinase reaction conditions were the same as those used with 2N4R Tau. SGK1- and GSK-3ß-mediated phosphorylation on 2N4R tau were examined using Phos-tag SDS-PAGE (Wako, Japan) (Kinoshita et al., 2009).

For a kinase assay following immunoprecipitation, SGK1, GSK-3ß and AKT were immunoprecipitated from brain homogenate or cell lines using dyna-beads protein A following the manufacturer’s instructions. The kinase activity of the immunoprecipitated hippocampal SGK1 was measured using an ADP-Glo kinase assay kit, using recombinant active SGK1 as a positive control following the manufacturer’s instructions (Promega). The kinase and ATPase reactions of SGK1 were carried out at 30°C for 1 hr using 400 μM ATP and a designed peptide (Murray et al., 2005) or recombinant tau as a substrate. EMD638683 (Chem Scene LLC, USA) was used as an SGK1 inhibitor. Luminescence was recoded as an integration of 1 sec using Mithras^2^ LB 943 Monochromator Multimode Microplate Reader (Berthold Technologies, Germany). The data were analyzed against recombinant SGK1, for which the unit activity was determined, and expressed as the amount of active SGK1 present per mg of brain lysate.

### Tubulin polymerization, tau aggregation and ThS assay

Recombinant tau from bacteria or heat-stable tau fractions isolated from mouse brains or SH-SY5Y cells stably expressing tau were incubated with 30 μM porcine brain tubulin (Cytoskeleton, Denver, CO, USA) in PEM buffer (80 mM PIPES, pH 6.9, 0.5 mM EGTA and 2 mM MgCl2) containing 0.5 mM guanidine triphosphate, 1 mM DTT and 15% glycerol at 37°C as described by the supplier. The assembly of tubulin into microtubules was monitored over time by a change in turbidity in a half area 96-well plate at 350 nm using SpectraMax iD3 Multi-Mode Microplate Readers (Molecular Devices). The absorbances were standardized and smoothed against the absorbance at the starting point.

Tau aggregates were prepared from purified recombinant tau protein. Tau (15 μM) and heparin (0.1 mg/ml) were incubated at 37°C for 72 h in 100 μl of 30 mM Tris-HCl, pH 7.5, containing 0.1% sodium azide and 20 mM DTT. In a time-dependent tau aggregation formation study, the amount of tau aggregates was monitored by ThS by mixing 10 μl of tau aggregate with 200 μl of 20 mM MOPS, pH 6.8, containing 5 μM ThS (Sigma). ThS fluorescence was measured using FlexStation II (Molecular Devices) (set at 436 nm excitation/535 nm emission) (Friedhoff et al., 1998). Transmission electron micrograph (TEM) images of the tau fibrils present in heparin-induced tau aggregates were captured by placing the fibrils on collodion-coated 300-mesh copper grids and stained with 2% (v/v) phosphotungstate using a HT7700 electron microscope (HITACHI).

### EMD638683 treatment

SGK1 inhibitor EMD638683, purchased from Chem Scene LLC (USA), was dissolved in DMSO to prepare stock solutions and kept at −80°C for further use. To confirm the kinase inhibitory specificity on tau phosphorylation, different doses of EMD638683 were added to the tau kinase reactions in the presence of SGK1 or GSK-3β. The inhibitory effect of EMD638683 was evaluated either by the ADP-Glo kinase assay or by Western blotting analysis using PS214 and PS396 anti-tau antibodies. For the cell-based study, SH-SY5Y cells expressing stable 2N4R tau were maintained in DMEM/F-12 medium (Sigma-Aldrich) supplemented with G418 and were allowed to grow in 6-well plates. After confirming 70% confluency, EMD638683 was applied at different concentrations and exposed for 24 hrs for dose-dependent analysis. The time-dependent study was carried out by using 1 μM EMD638683. An equal volume of DMSO was used as a vehicle. Treatment in the presence of DXM or hyperglycemic conditions was carried out by transferring the cells to a minimal medium containing 5.5 mM glucose and 2% Serum replacement (Gibco) before forty-eight hours of experiments. DXM, glucose and EMD638683 were applied accordingly and incubated for 6 hrs. For in vivo effect of EMD638683 on tau phosphorylation, 5.0 mg/kg EMD638683 were injected into the mouse brain hippocampus through stereotactic surgery. After 72 hrs, mice were sacrificed, and the effect were analyzed biochemically.

### Statistical analysis

JMP 12.0 software (SAS Institute, North Carolina, USA) was used to perform statistical analyses. Two-way factorial ANOVA followed by Tukey’s HSD was used for four-group comparisons. Mann-Whitney U test or Wilcoxon test were used when the distribution was not normal. For two-group comparisons with normal distribution, data were analyzed by Student’s *t*-test. In all cases, significance was noted at *p* < 0.05. Values are expressed as the means ± SEM. Blinding was performed in some key experiments (Immunohistochemical analysis for PS214 and PHF1 tau [Fig 2C-D, Figure 2 Supplement 1B-C], quantitative immunofluorescence analysis [Figure 5-figure supplement 1B, D, Figure 6F, and Figure 6-figure supplement 1C]).

### Funding

The Department of Diagnosis, Prevention and Treatment of Dementia is an endowed department in the graduate school of medicine and is supported by Eisai pharmaceuticals and Nihon Medi-Physics co., Japan. Montasir Elahi is supported by the Research Center for Old Age grant, Project-Kenkyo grant and Sportology Center grant, Juntendo University.

## Acknowledgments

Plasmid vectors for full length and CTF tau were provided by Dr. Shin-Ei Matsumoto. Authors thankful to the graduate students of the Department of Diagnosis, Prevention and Treatment of Dementia, Mohammed Nasir Uddin for his occasional help with reagent preparation.

## Authors Contribution

M.E. designed research, performed experiments, and analyzed data; S.S. and Y.I. occasionally help with the experiments. Y.M. and Y.I. supervised the whole study. H.H., M.T., K.I. and N.H. contributed new reagents/analytic tools; M.E., and Y.I. wrote the paper.

## Availability of the data, methods and materials

All gene expression or microarray data will be deposited in public domains and the detailed methods and reagents will be provided upon request.

## References

Ackermann, T.F., Boini, K.M., Beier, N., Scholz, W., Fuchss, T., and Lang, F. (2011). EMD638683, a novel SGK inhibitor with antihypertensive potency. Cell Physiol Biochem 28, 137–146.

Arnold, S.E., Arvanitakis, Z., Macauley-Rambach, S.L., Koenig, A.M., Wang, H.Y., Ahima, R.S., Craft, S., Gandy, S., Buettner, C., Stoeckel, L.E., et al. (2018). Brain insulin resistance in type 2 diabetes and Alzheimer disease: concepts and conundrums. Nat Rev Neurol 14, 168–181.

Bhat, N.R., and Thirumangalakudi, L. (2013). Increased tau phosphorylation and impaired brain insulin/IGF signaling in mice fed a high fat/high cholesterol diet. J Alzheimers Dis 36, 781–789.

Biessels, G.J., Staekenborg, S., Brunner, E., Brayne, C., and Scheltens, P. (2006). Risk of dementia in diabetes mellitus: a systematic review. Lancet Neurol 5, 64–74.

Blázquez, E., Velázquez, E., Hurtado-Carneiro, V., and Ruiz-Albusac, J.M. (2014). Insulin in the brain: its pathophysiological implications for States related with central insulin resistance, type 2 diabetes and Alzheimer’s disease. Front Endocrinol (Lausanne) 5, 161–161.

Cavallini, A., Brewerton, S., Bell, A., Sargent, S., Glover, S., Hardy, C., Moore, R., Calley, J., Ramachandran, D., Poidinger, M., et al. (2013). An unbiased approach to identifying tau kinases that phosphorylate tau at sites associated with Alzheimer disease. J Biol Chem 288, 23331–23347.

Chen, W., Chen, Y., Xu, B.-e., Juang, Y.-C., Stippec, S., Zhao, Y., and Cobb, M.H. (2009). Regulation of a third conserved phosphorylation site in SGK1. J Biol Chem 284, 3453–3460.

Chun, J., Kwon, T., Lee, E.J., Kim, C.H., Han, Y.S., Hong, S.K., Hyun, S., and Kang, S.S. (2004). 14-3-3 Protein mediates phosphorylation of microtubule-associated protein tau by serum- and glucocorticoid-induced protein kinase 1. Mol Cells 18, 360–368.

Cole, A., Frame, S., and Cohen, P. (2004). Further evidence that the tyrosine phosphorylation of glycogen synthase kinase-3 (GSK3) in mammalian cells is an autophosphorylation event. Biochem J 377, 249–255.

Dey, A., Hao, S., Wosiski-Kuhn, M., and Stranahan, A.M. (2017). Glucocorticoid-mediated activation of GSK3beta promotes tau phosphorylation and impairs memory in type 2 diabetes. Neurobiol Aging 57, 75–83.

Di, J., Cohen, L.S., Corbo, C.P., Phillips, G.R., El Idrissi, A., and Alonso, A.D. (2016). Abnormal tau induces cognitive impairment through two different mechanisms: synaptic dysfunction and neuronal loss. Sci Rep 6, 20833.

Elahi, M., Hasan, Z., Motoi, Y., Matsumoto, S.E., Ishiguro, K., and Hattori, N. (2016a). Region-Specific Vulnerability to Oxidative Stress, Neuroinflammation, and Tau Hyperphosphorylation in Experimental Diabetes Mellitus Mice. J Alzheimers Dis 51, 1209–1224.

Elahi, M., Motoi, Y., Matsumoto, S.E., Hasan, Z., Ishiguro, K., and Hattori, N. (2016b). Short-term treadmill exercise increased tau insolubility and neuroinflammation in tauopathy model mice. Neurosci Lett 610, 207–212.

Friedhoff, P., Schneider, A., Mandelkow, E.M., and Mandelkow, E. (1998). Rapid assembly of Alzheimer-like paired helical filaments from microtubule-associated protein tau monitored by fluorescence in solution. Biochemistry 37, 10223–10230.

Gan, W., Ren, J., Li, T., Lv, S., Li, C., Liu, Z., and Yang, M. (2018). The SGK1 inhibitor EMD638683, prevents Angiotensin II-induced cardiac inflammation and fibrosis by blocking NLRP3 inflammasome activation. Biochim Biophys Acta Mol Basis Dis 1864, 1–10.

GBD 2016 Dementia Collaborators (2019). Global, regional, and national burden of Alzheimer’s disease and other dementias, 1990-2016: a systematic analysis for the Global Burden of Disease Study 2016. Lancet Neurol 18, 88–106.

Geer, E.B., Islam, J., and Buettner, C. (2014). Mechanisms of glucocorticoid-induced insulin resistance: focus on adipose tissue function and lipid metabolism. Endocrinol Metab Clin North Am 43, 75–102.

Gratuze, M., El Khoury, N.B., Turgeon, A., Julien, C., Marcouiller, F., Morin, F., Whittington, R.A., Marette, A., Calon, F., and Planel, E. (2017). Tau hyperphosphorylation in the brain of ob/ob mice is due to hypothermia: Importance of thermoregulation in linking diabetes and Alzheimer’s disease. Neurobiol Dis 98, 1–8.

Green, K.N., Billings, L.M., Roozendaal, B., McGaugh, J.L., and LaFerla, F.M. (2006). Glucocorticoids increase amyloid-beta and tau pathology in a mouse model of Alzheimer’s disease. J Neurosci 26, 9047–9056.

Grundke-Iqbal, I., Iqbal, K., Tung, Y.C., Quinlan, M., Wisniewski, H.M., and Binder, L.I. (1986). Abnormal phosphorylation of the microtubule-associated protein tau (tau) in Alzheimer cytoskeletal pathology. Proc Natl Acad Sci U S A 83, 4913–4917.

Hanger, D.P., Lau, D.H., Phillips, E.C., Bondulich, M.K., Guo, T., Woodward, B.W., Pooler, A.M., and Noble, W. (2014). Intracellular and extracellular roles for tau in neurodegenerative disease. J Alzheimers Dis 40 Suppl 1, S37–45.

Hanger, D.P., and Noble, W. (2011). Functional implications of glycogen synthase kinase-3-mediated tau phosphorylation. Int J Alzheimers Dis 2011, 352805.

Hara, Y., Motoi, Y., Hikishima, K., Mizuma, H., Onoe, H., Matsumoto, S.E., Elahi, M., Okano, H., Aoki, S., and Hattori, N. (2017). Involvement of the Septo-Hippocampal Cholinergic Pathway in Association with Septal Acetylcholinesterase Upregulation in a Mouse Model of Tauopathy. Curr Alzheimer Res 14, 94–103.

Hasegawa, M., Crowther, R.A., Jakes, R., and Goedert, M. (1997). Alzheimer-like changes in microtubule-associated protein Tau induced by sulfated glycosaminoglycans. Inhibition of microtubule binding, stimulation of phosphorylation, and filament assembly depend on the degree of sulfation. J Biol Chem 272, 33118–33124.

Hinds, L.R., Chun, L.E., Woodruff, E.R., Christensen, J.A., Hartsock, M.J., and Spencer, R.L. (2017). Dynamic glucocorticoid-dependent regulation of Sgk1 expression in oligodendrocytes of adult male rat brain by acute stress and time of day. PLoS One 12, e0175075.

Iqbal, K., Liu, F., and Gong, C.-X. (2014). Alzheimer disease therapeutics: focus on the disease and not just plaques and tangles. Biochem Pharmacol 88, 631–639.

Iqbal, K., Liu, F., Gong, C.X., Alonso Adel, C., and Grundke-Iqbal, I. (2009). Mechanisms of tau-induced neurodegeneration. Acta Neuropathol 118, 53–69.

Ishiguro, K., Sato, K., Takamatsu, M., Park, J., Uchida, T., and Imahori, K. (1995). Analysis of phosphorylation of tau with antibodies specific for phosphorylation sites. Neurosci Lett 202, 81–84.

Johnson, L.A., Zuloaga, K.L., Kugelman, T.L., Mader, K.S., Morre, J.T., Zuloaga, D.G., Weber, S., Marzulla, T., Mulford, A., Button, D., et al. (2016). Amelioration of Metabolic Syndrome-Associated Cognitive Impairments in Mice via a Reduction in Dietary Fat Content or Infusion of Non-Diabetic Plasma. EBioMedicine 3, 26–42.

Kambe, T., Motoi, Y., Inoue, R., Kojima, N., Tada, N., Kimura, T., Sahara, N., Yamashita, S., Mizoroki, T., Takashima, A., et al. (2011). Differential regional distribution of phosphorylated tau and synapse loss in the nucleus accumbens in tauopathy model mice. Neurobiol Dis 42, 404–414.

Kim, J.Y., Lee, H.J., Lee, S.J., Jung, Y.H., Yoo, D.Y., Hwang, I.K., Seong, J.K., Ryu, J.M., and Han, H.J. (2017). Palmitic Acid-BSA enhances Amyloid-beta production through GPR40-mediated dual pathways in neuronal cells: Involvement of the Akt/mTOR/HIF-1alpha and Akt/NF-kappaB pathways. Sci Rep 7, 4335.

Kinoshita, E., Kinoshita-Kikuta, E., and Koike, T. (2009). Separation and detection of large phosphoproteins using Phos-tag SDS-PAGE. Nature Protocols 4, 1513.

Koga, S., Kojima, A., Kuwabara, S., and Yoshiyama, Y. (2014). Immunohistochemical analysis of tau phosphorylation and astroglial activation with enhanced leptin receptor expression in diet-induced obesity mouse hippocampus. Neurosci Lett 571, 11–16.

Kothari, V., Luo, Y., Tornabene, T., O’Neill, A.M., Greene, M.W., Geetha, T., and Babu, J.R. (2017). High fat diet induces brain insulin resistance and cognitive impairment in mice. Biochimica et Biophysica Acta (BBA) - Molecular Basis of Disease 1863, 499–508.

Krishnankutty, A., Kimura, T., Saito, T., Aoyagi, K., Asada, A., Takahashi, S.-I., Ando, K., Ohara-Imaizumi, M., Ishiguro, K., and Hisanaga, S.-i. (2017). In vivo regulation of glycogen synthase kinase 3β activity in neurons and brains. Sci Rep 7, 8602.

Lang, F., Bohmer, C., Palmada, M., Seebohm, G., Strutz-Seebohm, N., and Vallon, V. (2006). (Patho)physiological significance of the serum- and glucocorticoid-inducible kinase isoforms. Physiol Rev 86, 1151–1178.

Leboucher, A., Laurent, C., Fernandez-Gomez, F.J., Burnouf, S., Troquier, L., Eddarkaoui, S., Demeyer, D., Caillierez, R., Zommer, N., Vallez, E., et al. (2013). Detrimental effects of diet-induced obesity on tau pathology are independent of insulin resistance in tau transgenic mice. Diabetes 62, 1681–1688.

Li, P., Hao, Y., Pan, F.H., Zhang, M., Ma, J.Q., and Zhu, D.L. (2016). SGK1 inhibitor reverses hyperglycemia partly through decreasing glucose absorption. J Mol Endocrinol 56, 301–309.

Li, X.L., Aou, S., Oomura, Y., Hori, N., Fukunaga, K., and Hori, T. (2002). Impairment of long-term potentiation and spatial memory in leptin receptor-deficient rodents. Neuroscience 113, 607–615.

Liu, C.-C., Kanekiyo, T., Xu, H., and Bu, G. (2013). Correction: Apolipoprotein E and Alzheimer disease: risk, mechanisms and therapy. Nature Reviews Neurology 9, 184–184.

Liu, F., Li, B., Tung, E.J., Grundke-Iqbal, I., Iqbal, K., and Gong, C.X. (2007). Site-specific effects of tau phosphorylation on its microtubule assembly activity and self-aggregation. Eur J Neurosci 26, 3429–3436.

Liu, S.J., Zhang, J.Y., Li, H.L., Fang, Z.Y., Wang, Q., Deng, H.M., Gong, C.X., Grundke-Iqbal, I., Iqbal, K., and Wang, J.Z. (2004). Tau becomes a more favorable substrate for GSK-3 when it is prephosphorylated by PKA in rat brain. J Biol Chem 279, 50078–50088.

Livingston, G., Sommerlad, A., Orgeta, V., Costafreda, S.G., Huntley, J., Ames, D., Ballard, C., Banerjee, S., Burns, A., Cohen-Mansfield, J., et al. (2017). Dementia prevention, intervention, and care. Lancet 390, 2673–2734.

Livingstone, D.E.W., Grassick, S.L., Currie, G.L., Walker, B.R., and Andrew, R. (2009). Dysregulation of glucocorticoid metabolism in murine obesity: comparable effects of leptin resistance and deficiency. J Endocrinol 201, 211–218.

Mason, B.L., Pariante, C.M., Jamel, S., and Thomas, S.A. (2010). Central nervous system (CNS) delivery of glucocorticoids is fine-tuned by saturable transporters at the blood-CNS barriers and nonbarrier regions. Endocrinology 151, 5294–5305.

Matsumoto, S.E., Motoi, Y., Ishiguro, K., Tabira, T., Kametani, F., Hasegawa, M., and Hattori, N. (2015). The twenty-four KDa C-terminal tau fragment increases with aging in tauopathy mice: implications of prion-like properties. Hum Mol Genet 24, 6403–6416.

McEwen, B.S. (1997). Possible mechanisms for atrophy of the human hippocampus. Mol Psychiatry 2, 255–262.

Morris, J.K., Vidoni, E.D., Wilkins, H.M., Archer, A.E., Burns, N.C., Karcher, R.T., Graves, R.S., Swerdlow, R.H., Thyfault, J.P., and Burns, J.M. (2016). Impaired fasting glucose is associated with increased regional cerebral amyloid. Neurobiol Aging 44, 138–142.

Mrak, R.E. (2009). Alzheimer-type neuropathological changes in morbidly obese elderly individuals. Clin Neuropathol 28, 40–45.

Mullins, R.J., Diehl, T.C., Chia, C.W., and Kapogiannis, D. (2017). Insulin Resistance as a Link between Amyloid-Beta and Tau Pathologies in Alzheimer’s Disease. Front Aging Neurosci 9, 118–118.

Murray, J.T., Cummings, L.A., Bloomberg, G.B., and Cohen, P. (2005). Identification of different specificity requirements between SGK1 and PKBα. FEBS Lett 579, 991–994.

Noble, W., Hanger, D.P., Miller, C.C., and Lovestone, S. (2013). The importance of tau phosphorylation for neurodegenerative diseases. Front Neurol 4, 83.

Park, J., Leong, M.L., Buse, P., Maiyar, A.C., Firestone, G.L., and Hemmings, B.A. (1999). Serum and glucocorticoid-inducible kinase (SGK) is a target of the PI 3-kinase-stimulated signaling pathway. Embo J 18, 3024–3033.

Peng, D., Pan, X., Cui, J., Ren, Y., and Zhang, J. (2013). Hyperphosphorylation of tau protein in hippocampus of central insulin-resistant rats is associated with cognitive impairment. Cell Physiol Biochem 32, 1417–1425.

Planel, E., Tatebayashi, Y., Miyasaka, T., Liu, L., Wang, L., Herman, M., Yu, W.H., Luchsinger, J.A., Wadzinski, B., Duff, K.E., et al. (2007). Insulin dysfunction induces in vivo tau hyperphosphorylation through distinct mechanisms. J Neurosci 27, 13635–13648.

Platt, T.L., Beckett, T.L., Kohler, K., Niedowicz, D.M., and Murphy, M.P. (2016). Obesity, diabetes, and leptin resistance promote tau pathology in a mouse model of disease. Neuroscience 315, 162–174.

Revsin, Y., Rekers, N.V., Louwe, M.C., Saravia, F.E., De Nicola, A.F., de Kloet, E.R., and Oitzl, M.S. (2008). Glucocorticoid Receptor Blockade Normalizes Hippocampal Alterations and Cognitive Impairment in Streptozotocin-Induced Type 1 Diabetes Mice. Neuropsychopharmacology 34, 747.

Sadik, G., Tanaka, T., Kato, K., Yamamori, H., Nessa, B.N., Morihara, T., and Takeda, M. (2009). Phosphorylation of tau at Ser214 mediates its interaction with 14-3-3 protein: implications for the mechanism of tau aggregation. J Neurochem 108, 33–43.

Sahin, P., McCaig, C., Jeevahan, J., Murray, J.T., and Hainsworth, A.H. (2013). The cell survival kinase SGK1 and its targets FOXO3a and NDRG1 in aged human brain. Neuropathol Appl Neurobiol 39, 623–633.

Sajan, M., Hansen, B., Ivey, R., 3rd, Sajan, J., Ari, C., Song, S., Braun, U., Leitges, M., Farese-Higgs, M., and Farese, R.V. (2016). Brain Insulin Signaling Is Increased in Insulin-Resistant States and Decreases in FOXOs and PGC-1alpha and Increases in Abeta1-40/42 and Phospho-Tau May Abet Alzheimer Development. Diabetes 65, 1892–1903.

Schwab, M., Lupescu, A., Mota, M., Mota, E., Frey, A., Simon, P., Mertens, P.R., Floege, J., Luft, F., Asante-Poku, S., et al. (2008). Association of SGK1 gene polymorphisms with type 2 diabetes. Cell Physiol Biochem 21, 151–160.

Sohn, J., Takahashi, M., Okamoto, S., Ishida, Y., Furuta, T., and Hioki, H. (2017). A Single Vector Platform for High-Level Gene Transduction of Central Neurons: Adeno-Associated Virus Vector Equipped with the Tet-Off System. PLoS One 12, e0169611.

Spauwen, P.J., Kohler, S., Verhey, F.R., Stehouwer, C.D., and van Boxtel, M.P. (2013). Effects of type 2 diabetes on 12-year cognitive change: results from the Maastricht Aging Study. Diabetes Care 36, 1554–1561.

Stieler, J.T., Bullmann, T., Kohl, F., Tøien, Ø., Brückner, M.K., Härtig, W., Barnes, B.M., and Arendt, T. (2011). The Physiological Link between Metabolic Rate Depression and Tau Phosphorylation in Mammalian Hibernation. PLoS One 6, e14530.

Stranahan, A.M., Arumugam, T.V., Cutler, R.G., Lee, K., Egan, J.M., and Mattson, M.P. (2008). Diabetes impairs hippocampal function through glucocorticoid-mediated effects on new and mature neurons. Nat Neurosci 11, 309–317.

Takalo, M., Haapasalo, A., Martiskainen, H., Kurkinen, K.M., Koivisto, H., Miettinen, P., Khandelwal, V.K., Kemppainen, S., Kaminska, D., Makinen, P., et al. (2014). High-fat diet increases tau expression in the brain of T2DM and AD mice independently of peripheral metabolic status. J Nutr Biochem 25, 634–641.

Ullrich, S., Berchtold, S., Ranta, F., Seebohm, G., Henke, G., Lupescu, A., Mack, A.F., Chao, C.M., Su, J., Nitschke, R., et al. (2005). Serum- and glucocorticoid-inducible kinase 1 (SGK1) mediates glucocorticoid-induced inhibition of insulin secretion. Diabetes 54, 1090–1099.

Valente, T., Gella, A., Fernandez-Busquets, X., Unzeta, M., and Durany, N. (2010). Immunohistochemical analysis of human brain suggests pathological synergism of Alzheimer’s disease and diabetes mellitus. Neurobiol Dis 37, 67–76.

Virdee, K., Yoshida, H., Peak-Chew, S., and Goedert, M. (2007). Phosphorylation of human microtubule-associated protein tau by protein kinases of the AGC subfamily. FEBS Lett 581, 2657–2662.

Webster, M.K., Goya, L., Ge, Y., Maiyar, A.C., and Firestone, G.L. (1993). Characterization of sgk, a novel member of the serine/threonine protein kinase gene family which is transcriptionally induced by glucocorticoids and serum. Mol Cell Biol 13, 2031–2040.

Webster, S.J., Bachstetter, A.D., and Van Eldik, L.J. (2013). Comprehensive behavioral characterization of an APP/PS-1 double knock-in mouse model of Alzheimer’s disease. Alzheimers Res Ther 5, 28.

Yang, Y.C., Lin, C.H., and Lee, E.H. (2006). Serum- and glucocorticoid-inducible kinase 1 (SGK1) increases neurite formation through microtubule depolymerization by SGK1 and by SGK1 phosphorylation of tau. Mol Cell Biol 26, 8357–8370.

